# Hidden Knowledge Recovery from GAN-generated Single-cell RNA-seq Data

**DOI:** 10.1101/2023.11.27.568840

**Authors:** Najeebullah Shah, Fanhong Li, Xuegong Zhang

## Abstract

**Background:** Machine learning methods have recently been shown powerful in discovering knowledge from scientific data, offering promising prospects for discovery learning. In the meanwhile, Deep Generative Models like Generative Adversarial Networks (GANs) have excelled in generating synthetic data close to real data. GANs have been extensively employed, primarily motivated by generating synthetic data for privacy preservation, data augmentation, etc. However, certain dimensions of GANs have received limited exploration in current literature. Existing studies predominantly utilize huge datasets, presenting a challenge when dealing with limited, complex datasets. Researchers have highlighted the ineffectiveness of conventional scores for selecting optimal GANs on limited datasets that exhibit complex high order relationships. Furthermore, current methods evaluate GAN’s performance by comparing synthetic data to real data without assessing the preservation of high-order relationships. Researchers have advocated for more objective GAN evaluation techniques and emphasized the importance of establishing interpretable connections between GAN latent space variables and meaningful data semantics.

**Results:** In this study, we used a custom GAN model to generate quality synthetic data for a very limited, complex biological dataset. We successfully recovered cell-lineage developmental story from synthetic data using the ab-initio knowledge discovery method, we previously developed. Our custom GAN model performed better than state-of-the-art cscGAN model, when evaluated for recovering hidden knowledge from limited, complex dataset. Then we devise a temporal dataset specific quantitative scoring mechanism to successfully reproduce GAN results for human and mouse embryonic datasets. Our Latent Space Interpretation (LSI) scheme was able to identify anomalies. We also found that the latent space in GAN effectively captured the semantic information and may be used to interpolate data when the sampling of real data is sparse.

**Conclusion:** In summary we used a customized GAN model to generate synthetic data for limited, complex dataset and compared the results with state-of-the-art cscGAN model. Cell-lineage developmental story is recovered as hidden knowledge to evaluate GAN for preserving complex high-order relationships. We formulated a quantitative score to successfully reproduce results on human and mouse embryonic datasets. We designed a LSI scheme to identify anomalies and understand the mechanism by which GAN captures important data semantics in its latent space.

## 1 Background

Machine Learning (ML) methods have made significant advances, especially for pattern recognition tasks, in making precise predictions and recognizing detailed patterns. Moreover, various efforts have been made to utilize ML models for discovering knowledge from scientific data [1–3]. Previously, we developed ML-based algorithm to discover scientific knowledge from real biological data in *ab-initio* manner [1]. The recovered knowledge was remarkably similar to existing knowledge, indicating the potential of the ML-based method for discovering knowledge from new data.

Meanwhile, DGMs have been widely used in a broad range of research fields for a variety of applications [4–7]. The majority of research and applications are either directly or indirectly related to computer vision and image processing [8, 9]. In the field of medical sciences and health-care, DGMs are predominantly used in medical imaging [10, 11]. Some of the prominent applications of DGMs in bioinformatics are creating artificial human genome [12], cell gene imputation [13], generating protein structures [14], scRNA-seq dimentionality reduction [15], encouraging disentanglement [16] and simulating realistic scRNA-seq data [17, 18].

DGMs, especially GANs [19], have made significant advances for rapidly closing gap between real and synthetic samples. GANs have been extensively employed to generate valuable synthetic data in different scenarios. These scenarios include generating hard-to-obtain data from easy-to-obtain data [20], acquiring expensive MRI scans from relatively cheaper CT scans [21], augmenting synthetic scRNA-seq samples of specific cell types to counter bias and imbalance in datasets [18] and produce synthetic samples to protect data privacy [22]. In the context of Canada’s response to the COVID-19 pandemic, a notable challenge has emerged due to the implementation of the “privacy chill” policy, preventing the sharing of crucial health data. Experts have recognized the adverse effects of this policy on the country’s pandemic response efforts. To address this issue, DGM-generated synthetic data has been proposed as a viable solution [23]. Such synthetic datasets possesses the unique characteristic of anonymizing the underlying information, thereby addressing the privacy concerns associated with health data sharing. However, researchers have articulated the need to channel research works towards specific dimensions of GANs that have relatively received limited attention. Existing studies predominantly employ substantial datasets, over-shadowing the serious challenges that can be encountered while training GANs on very small sampled datasets. Researchers have emphasized on demonstrating the applicability of DGMs in scenarios where the size of the dataset is limited [24–26]. In this regard, carefully crafted GANs could be the most effective and suitable tool. Within such constrained contexts, an intriguing avenue of research lies in discovery learning from synthetic data. In literature, evaluating the generative models for producing quality synthetic data is an extremely difficult task [18, 27–29]. Current evaluation techniques compare real and synthetic data without validating whether generated data preserved complex high-order relations in the real data. Researchers have advocated for a more objective evaluation of GANs. Additionally, conventional quantitative methods like inception score [30] etc proves ineffective when the dataset exhibits complex properties. Therefore it is often challenging to reproduce the results. There is a pressing need to formulate a more dataset specific quantitative scores for reproducing GAN results specially on small, complex datasets [29]. Furthermore, the interpretation of GAN’s latent space remains a challenge. The development of an easy-to-understand frameworks have been endorsed to comprehend the nuanced relationship between GAN’s latent space variables and meaningful data semantics [25, 31].

In biological studies, single-cell genomics data give researchers the opportunity to study gene expression profiles at the resolution of individual cells. Researchers have used single cell RNA sequencing (scRNA-seq) to study cell heterogeneity, compare healthy cells from diseased cells, cell progression over a period of time, etc. The study of early human embryonic cell development is an important application of scRNA-seq data. Early human embryonic development refers to development of embryo from 8 cell stage until pre-implantation, a complicated process controlled by precise cellular behaviors [32]. We used early human embryonic dataset of 1529 cells, sequenced in five time-points from day3 (E3) to day7 (E7). The cells go through developmental phase during these five time-points differentiating into various major cell-lineages [33]. The size of the dataset falls significantly below the typical sample size required to effectively train DGMs. As such, this dataset aligns exceptionally well with our research objective, which entails training GANs on limited dataset and subsequently comparing the outcomes against those achieved by the state-of-the-art cscGAN model. Additionally, to substantiate the effectiveness of our experimental setup, we employed a limited dataset comprising only 1724 samples for the analysis of early mouse embryonic scRNA-seq data [34].

We conducted a set of experiments to investigate the different objectives this research intends to address. The general overview of our research work is demonstrated in Fig 1. We introduced a customized GAN method named cwGAN by incorporating the ideas of Conditional GAN [35] and Wasserstein GAN [36] with Gradient Penalty using Label smoothing [37]. The cwGAN is used to synthesize a limited scRNA-seq dataset of early human embryonic cell development. Then, we compared the results of our customized cwGAN with state-of-the-art cscGAN model for the same dataset. After that, we formulated a quantitative score, Time-point T-PCAVR (Time-point PCA Variance Ratio) error, to reproduce results by automatically selecting the most optimal GAN hyper-parameters. Furthermore, we applied the *ab-initio* knowledge discovery method, we developed earlier [1], to evaluate whether GAN preserved complex high-order relationships as hidden knowledge. Additionally, we developed LSI schemes to investigate the ability of GANs for preserving hidden knowledge and identifying anomalies. Results show that the synthetic data generated by our customized GAN was superior than cscGAN model on limited dataset. T-PCAVR quantitative error score enables the reproduction of quality synthetic data, for human and mouse embryonic scRNA-seq datasets, summarized in sub-section 3.11 and 3.12. However, T-PCAVR score have a limitation, explained in sub-section 3.13. Utilizing the *ab-initio* knowledge discovery method, the cell developmental information is successfully recovered from both real and synthetic data as hidden knowledge. Additional experiments, with LSI schemes, show that GAN was able to preserve high-order relations by capturing cell developmental story as unknown semantic in the latent space. The designed Time-point Cell-Lineage LSI scheme identified anomalies with conflicting time-point and cell-lineage labels. Time-point Cell-lineage LSI scheme also provided a more precise sampling of desired time-point and cell-lineage with the help of a comprehensively generated dataset. The T-PCAVR quantitative score effectively determined the optimal hyper-parameters for the cwGAN model, enabling the recovery of hidden knowledge from the early mouse embryonic scRNA-seq dataset.

**Fig. 1:**
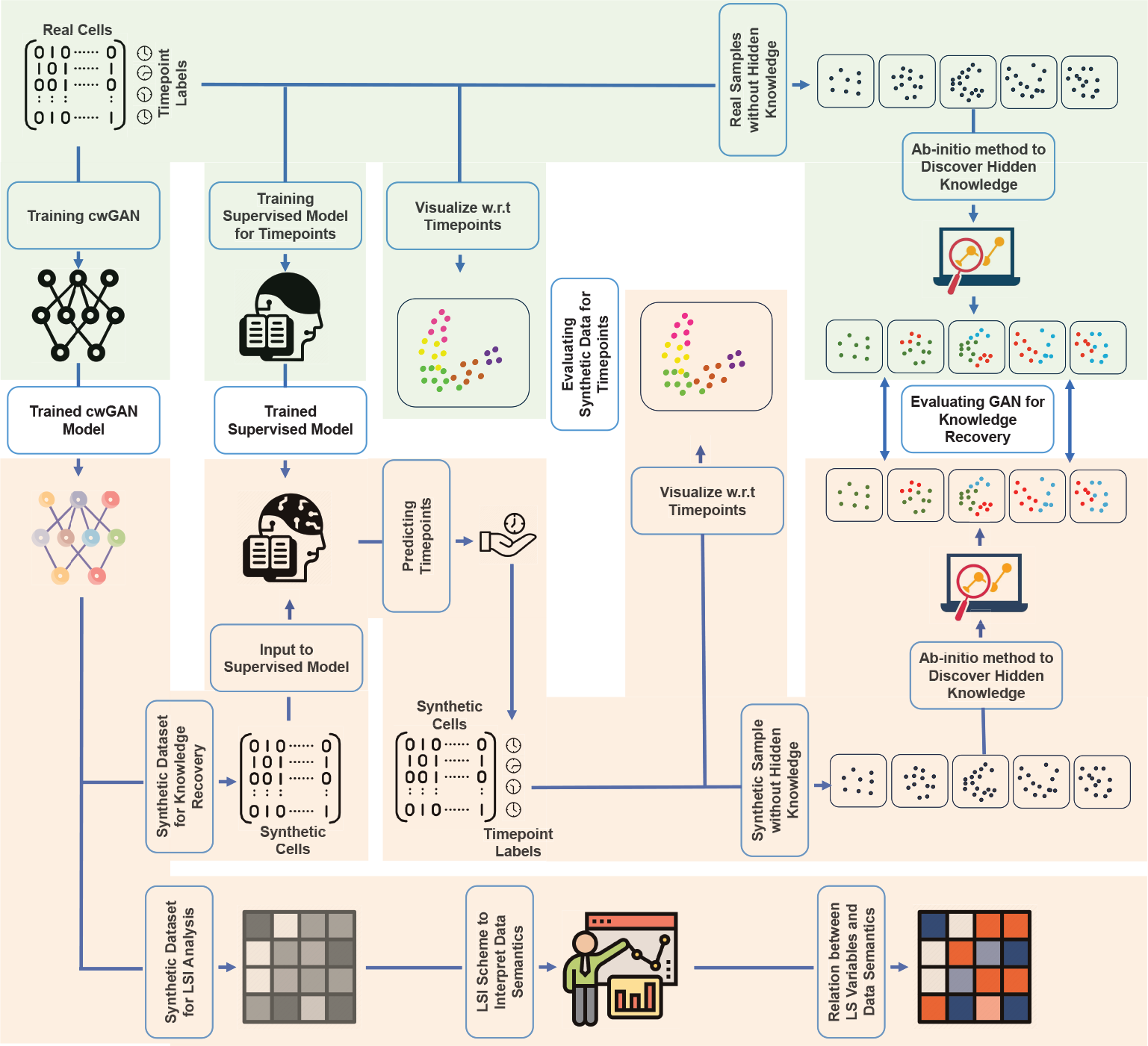
General Overview of this Research Work

## 2 Methods

### 2.1 GAN for Synthetic Data

We used GAN to generate synthetic data. Standard GAN, as shown in Fig. 2, did not perform well on the dataset we used in this work. Therefore, we made a number of changes to GAN model and make it better adapt to our problem. In this regard, we modified standard GAN by incorporating Wasserstein distance [36], an approximation of Earth Mover distance, with Gradient Penalty to avoid the mode collapse and instability during training. To mitigate the challenge of overfitting, we strategically employed time-points information as conditional input, integrating the concept of Conditional GAN [35]. Additionally, we used label smoothing [37] to introduce noise on conditional input as an additional regularization step. So finally we call this customized GAN as cwGAN by incorporating the ideas of Conditional GAN and Wasserstein GAN with Gradient Penalty using Label Smoothing. Results were further improved by reducing original counts of 26,178 genes to 490 highly differentiated genes. Generator loss is back-propagated once after every five iterations of back-propagating Discriminator loss.

**Fig. 2:**
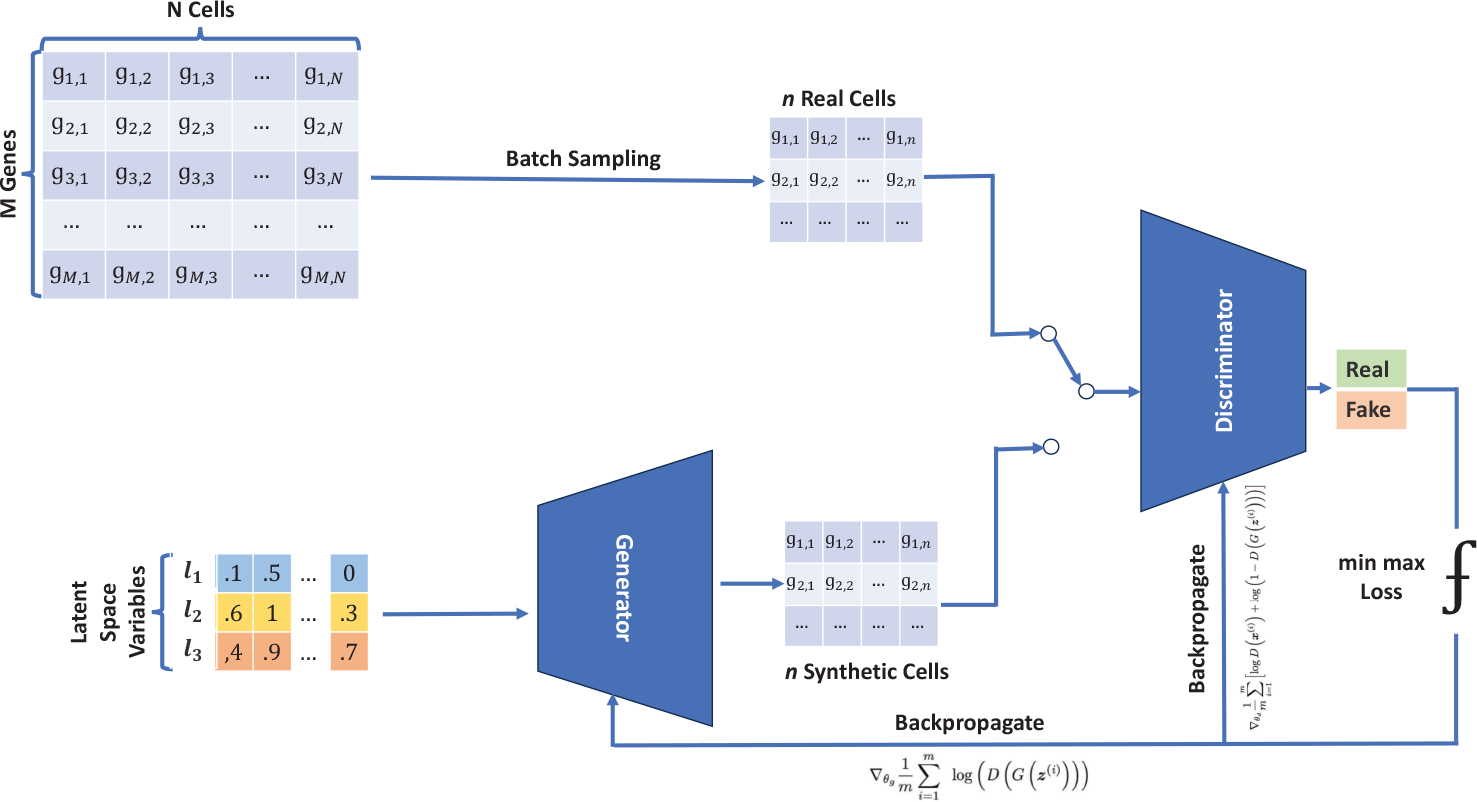
Standard GAN

Fig. 3 shows the conceptual structure of the customized GAN, we used to generate synthetic data for two main tasks. First, we generated synthetic dataset of 1529 samples against real dataset of 1529 samples. These datasets are used in recovering hidden knowledge from real and synthetic datasets for comparison. Second, we generated synthetic samples to find how GAN preserve hidden knowledge via LSI scheme.

**Fig. 3:**
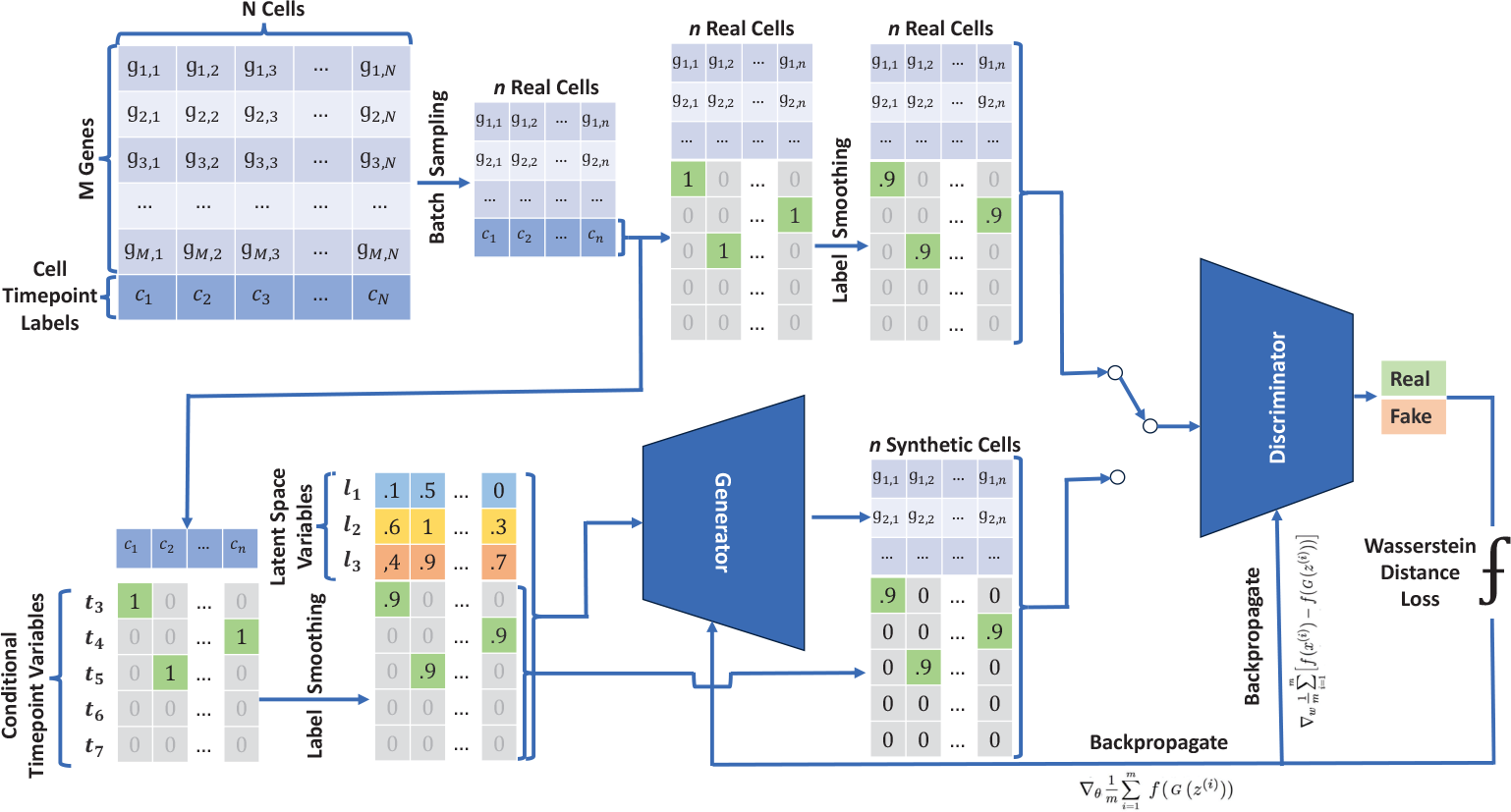
Structure of our Customized GAN

### 2.2 T-PCAVR Quantitative Score for Selecting Optimal GAN hyper-parameters

Initially, we considered using quantitative scores like loss values of Generator and Discriminator and a commonly used inception score [30] to automatically select the most optimal GAN model, during training. However, none of these quantitative scores were good indicators of optimally trained GAN. The ineffectiveness of these scores is also highlighted in literature [29]. Therefore we formulated a dataset specific score to select GAN with most optimal hyper-parameters. We utilized a cascaded approach of time-point and PCA variance ratio error scores. Time-point error is calculated as the absolute difference between the number of samples in real data and the number of samples in synthetic data for each time-point separately, divided by the number of real samples in that particular time-point. The error for each time-point accumulates to output the final Time-point Error Score. Algo. 1 shows the calculation of the total Time-point error.

Minimizing time-point errors is crucial for ensuring that synthetic data align with the real data in terms of sample numbers across different time points. Additionally, keeping a low PCAVR error score is vital to validate the inherent structure of the real data within the synthetic dataset. In literature, the variance explained by principal components have been utilized in an experimental setup to avoid dominant variation in a target dataset which might also be present in the background dataset [38]. Rather background dataset is used as reference to identify intricate interesting properties specific to the target dataset. In our experimental setup, we compare the percentage of variance explained by the first two principal components for samples of each time-point from synthetic and real datasets. Algo. 2 shows the calculation of the total PCAVR error.

#### Algorithm 1

Calculate Total Time-point Error

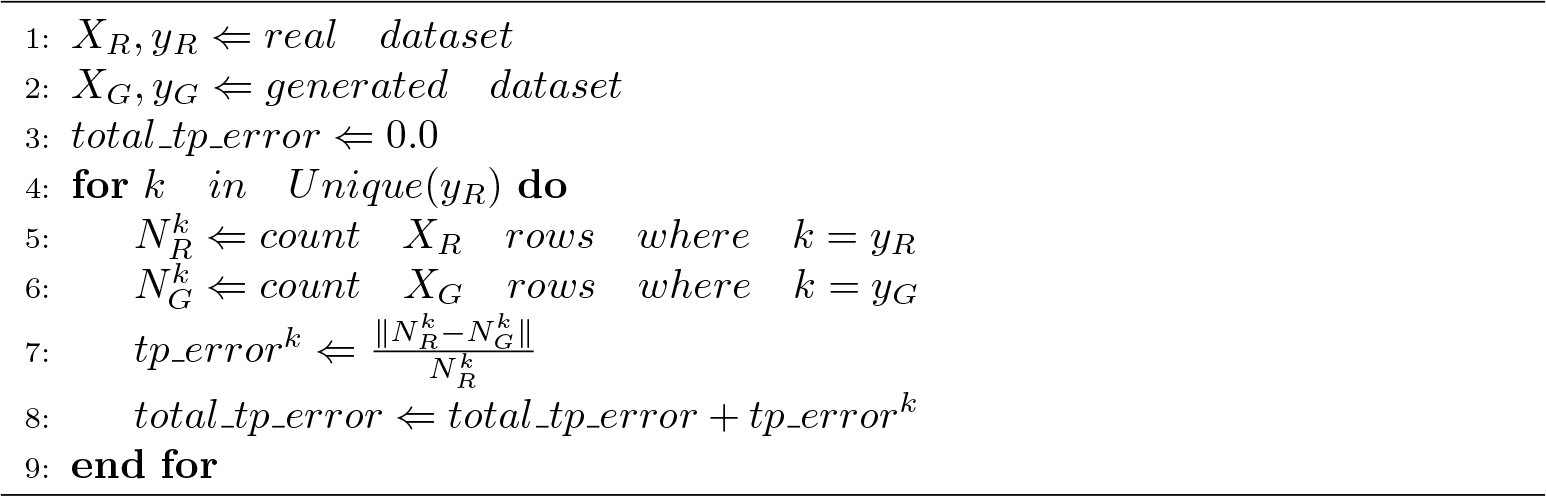

#### Algorithm 2

Calculate Total PCAVR Error

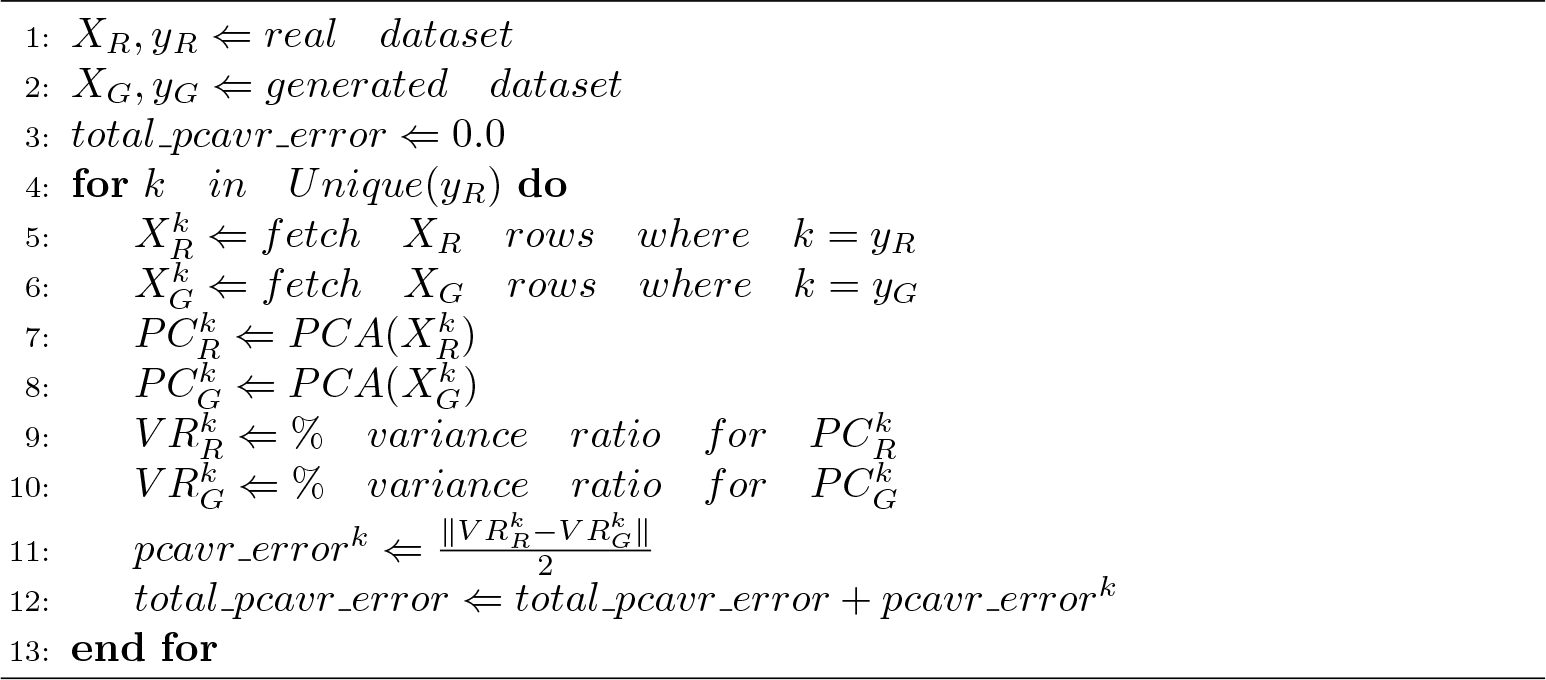

### 2.3 Supervised ML model for assigning time-points to synthetic data

Although we used time-point information as conditional input while generating synthetic dataset but we did not rely on conditional input to decide the time-point of the generated sample. Instead, we trained supervised ML model to learn the time-points of real data and then identified time-points of generated synthetic data. The reason for choosing identified time-points with supervised ML model over time-point information as conditional input is described in Results Section. To this end, we trained Support Vector Machine (SVM) model with Radial Basis Function (RBF) kernel [39] to learn time-points from real data. The trained SVM model is then used to assign time-points for the synthetic dataset. Both real and synthetic datasets are qualitatively assessed in terms of time-points using 2D principle component (PC) plots. The results are also quantitatively summarized in tabular form.

The supervised task of identifying five time-points for synthetic dataset is a multi-class classification problem. We adopt the OVR (One Vs Rest) strategy, which splits a multi-class classification into one binary classification per class, since SVM is designed for binary classification.

SVM did a good job learning time-points in the human embryonic dataset. But for the mouse embryonic dataset, SVM made a mistake by labeling all samples from different time-points as E6.5. So, we switched to Random Decision Forest (RDF) [40] for better accuracy. RDF learned the time-points from the real mouse data and accurately predicted them for the synthetic mouse data. We used 100 decision trees in RDF, each with a maximum depth of 10.

SVM did not perform well on the mouse embryonic dataset because the samples from different time-points were not clearly separated compared to human embryonic dataset. For a intuitive understanding, we used SVM with RBF kernel to plot the decision boundary for both human Fig. 4(left) and mouse Fig. 4(right) datasets. We simplified the data using PCA and then trained SVM. Upon inspection, we found that the decision boundary for mouse dataset was very poorly segregated compared to human dataset. While Fig. 4, utilizing two principal components as features, does not exactly delineate the decision boundary for the SVM with RBF kernel trained on a dataset comprising 490 features, it offers a valuable insight into the limitation of the SVM model in capturing temporal patterns within the early mouse embryonic dataset.

**Fig. 4:**
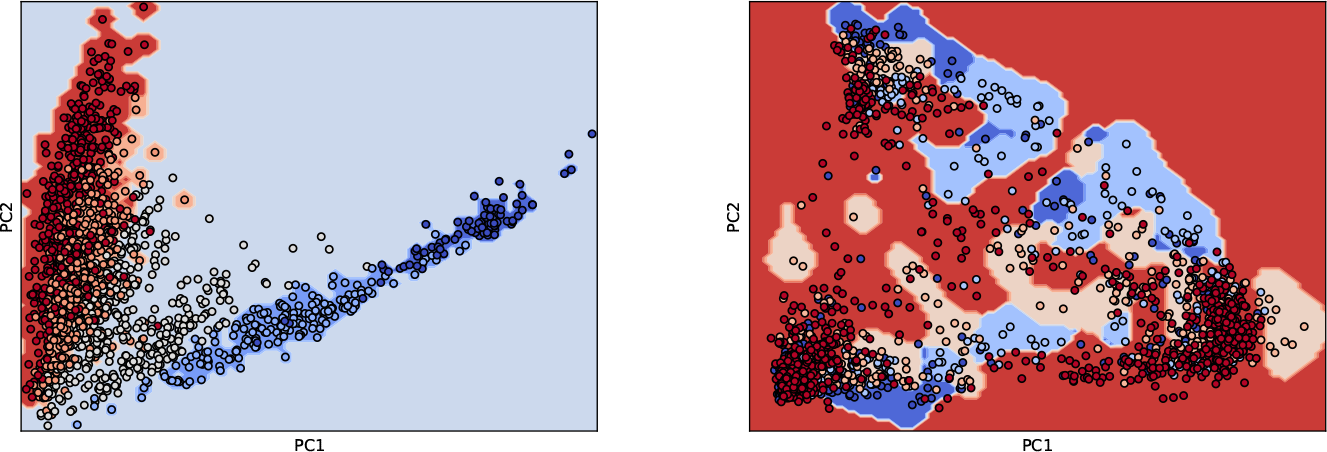
Decision boundary of SVM with RBF kernel for early human embryonic dataset (left) and early mouse embryonic dataset (right)

### 2.4 ab-intio Knowledge Discovery Method for Recovering Complex High Order Inter Relationships

*ab-initio* knowledge discovery method is the core algorithm of our previous research work. The method is a fully automated algorithm that quantifies the potential of samples from each time-point to become reference for creating cell-lineage developmental story [1]. The method presumes that one of the time-point has representative samples for all cell-lineages. We created 45 possible combinations with five time-points and cluster numbers in the range 2,…,10. The algorithm assigns Adjusted Reliability Score (ARS) to each combination. Samples of a time-point and cluster number combination that achieve maximum ARS is selected as reference. Samples of reference time-point is clustered with unsupervised learning model to get cell-lineage labels for each sample. A supervised model is trained to learn cell-lineages of reference time-point as target. The trained model is then used to identify cell-lineages for samples of remaining time-points. The method is individually applied to real and synthetic datasets. It is noteworthy to mention that in our study, we employed labels A, B, and C to denote cell-lineages, rather than using the actual names. This choice was deliberate, aimed at emphasizing that the acquisition of the cell-lineage developmental story was conducted in an ab initio manner. It is pertinent to note that in our prior research, a downstream analysis experiment was conducted to map these labels back to the actual cell-lineage names.

### 2.5 Assessment of the Recovered Hidden Knowledge

*ab-initio* knowledge discovery algorithm provides us with cell-lineage developmental story of real and synthetic datasets for every time-point. The two stories are expected to have significant resemblance in terms of cell-lineages in each time-point. This resemblance in cell-lineage developmental story serves as a critical evaluation criterion for GAN in preserving complex high order inter relationships. We used 2D PC plots for qualitative assessment of the two stories. Five 2D PC plots are used corresponding to five time-point in each story. Samples in each 2D PC plot are labeled for cell-lineages. The results are further summarized in tabular form for quantitative analysis. We also use stack plots for the combined, qualitative and quantitative, assessment of results.

### 2.6 Latent Space Interpretation Schemes

To investigate how GAN capture high-order relations, we study the encoding of important semantics in the latent space of GAN. GANs have been extremely successful at encoding important semantics about data in entirely unsupervised experimental setup [41]. It has been shown that several semantics emerge in the latent space of GAN during training [42]. The semantic encoding illustrates smooth and linear directions that affect properties of data (time-points and cell-lineages) w.r.t latent space variables [41]. Researchers have emphasized the importance of establishing interpretable connections between GAN’s latent space variables and meaningful data semantics. In this regard, we performed three LSI experiments to interpret time-points and cell-lineages w.r.t latent space variables. These LSI experiments are represented with a general schematic view shown in Fig. 5. The first two experiments help us understand the effect of small change in latent space variables on the corresponding time-points and cell-lineages of synthetic samples. These two experiments are used to investigate the ability of GAN for capturing relationship of time-points as known and cell-lineages as unknown semantic with the latent space variables. The successful capturing of known and unknown semantics will confirm that GAN preserved hidden knowledge (cell-lineages) in the same way as the known information (time-points). We then performed the LSI experiment, w.r.t time-points and cell-lineages combined, to gain deeper insights and identify anomalies within the synthetic data. In all of these experiments, time-point and cell-lineage are identified using supervised ML models. To visualize the impact of small alterations in latent space variables onto important data semantics, we decided to construct 2D Interpretation plots. Since we have 3 latent space variables to be represented on two-dimensional plots, defined along the x-y axis, therefore we create three possible sets of two latent variable values keeping one fixed at a time. These three sets are shown in Eq. 1, Eq. 2 and Eq. 3.

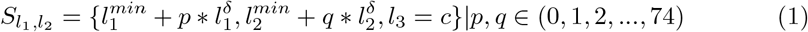

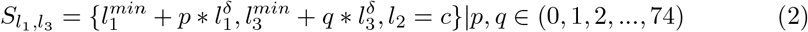

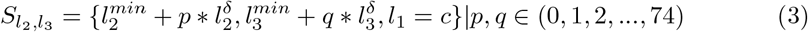

**Fig. 5:**
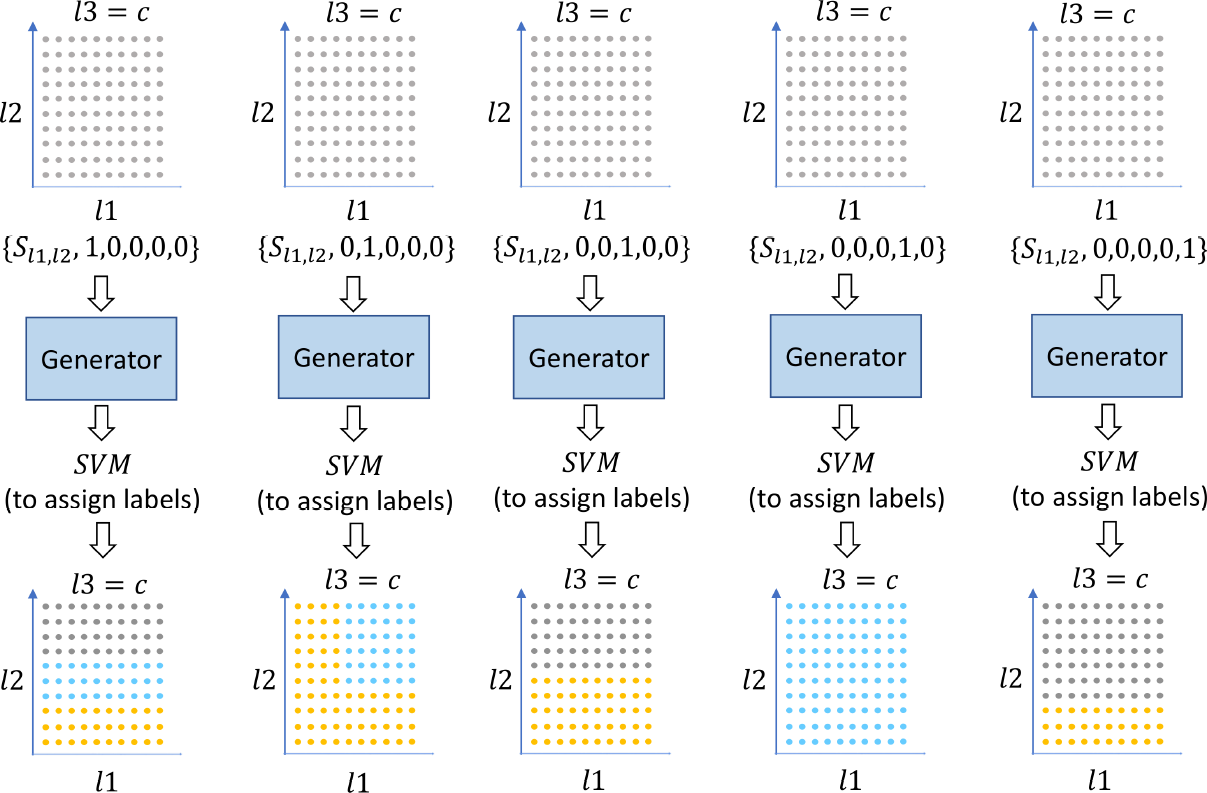
General Schematic View for Latent Space Interpretation

Where 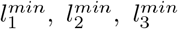 are the minimum values of latent space variables. We decided to use 75 values for each latent space variable as *p* and *q*, incremented with fix step size of 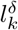 shown in Eq. 4. *c* represents constant value for one of the latent space variables in each 2D interpretation plot.

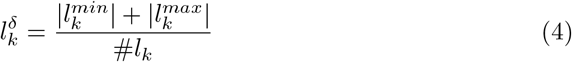

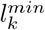 and 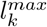 is minimum and maximum values for each latent space variable from the 1529 samples of synthetic dataset.

## 3 Results

### 3.1 The Dataset

In this study, we used early human embryonic cell development scRNA-seq dataset. The dataset consists of gene expression profile from 1529 individual cells. There are 490 highly differentiated genes in every cell. Each of 1529 cells are sequenced in one of five time-points i.e., day3 (E3) to day7 (E7). The number of cells in E3 to E7 are 81, 180, 377, 415, 466 respectively. These cells belong to different developmental stages. Time-points are known information and cell-lineages are hidden knowledge.

### 3.2 GAN Generates Quality Synthetic Data

GAN is used as deep generative model to find the underlying distribution of real 1529 cell dataset. Trained GAN Generator is used to generate 1529 samples of synthetic dataset. We used supervised model to learn time-points from real data. Then used trained supervised model to assign time-point to synthetic data. Fig. 6 show the projection of 490 genes onto 2D PC plots for real (left) and synthetic (right) datasets labelled with time-points. Same colors are used in both subplots to represent time-points of E3 (purple), E4 (orange), E5 (yellow), E6 (brown) and E7 (pink). These results are also summarized in Tab. 2

**Fig. 6:**
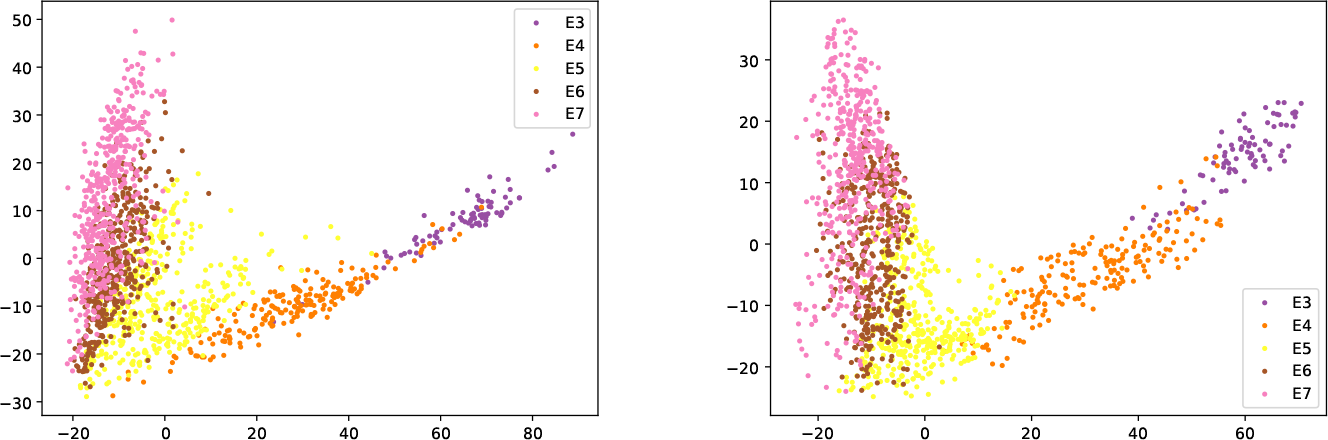
2d PC projection of 490 gene samples for real (left) and generated (right) datasets

**Table 1:** List of controlled parameter names and the set of values for each controlled parameter.

| Controlled Parameter Name | Controlled Parameter Values |
| --- | --- |
| Number of Latent Space Variables | 2, **3**, 5, 10, 20 |
| Number of Hidden Layers | 2, **3**, 4 |
| Number of Neurons in First Hidden Layer | 32, 64, 100, 128, **200**, 256, 300, 350, 400, 512, 1024 |
| Number of Neurons in Second Hidden Layer | 64, 100, 128, 200, 256, 300, **350**, 400, 500, 1024, 2048 |
| Number of Neurons in Third Hidden Layer (If applied) | 128, 200, 256, 300, 350, 400, **500**, 1024, 2048 |
| Number of Neurons in Fourth Hidden Layer (If applied) | 200, 256, 300, 350, 400, 500, 1024, 2048 |
| Number of Genes | **490**, 26, 178 |
| Neuron Initialization techniques | **Xavier**, He |
| Sampling Method for latent variables | **Uniform**, Normal |
| Batch Sizes | 32, **64**, 128 |
| Values for Label Smoothing | 0.95, 0.96, **0.97**, 0.98, 0.99, 1.00 |
| Optimizer Algorithm | **RMSProp**, Adam |
| Learning Rate for the loss function of Generator | 0.05, **0.005**, 0.0005 |
| Learning Rate for the loss function of Discriminator | 0.05, **0.005**, 0.0005 |

### 3.3 T-PCAVR Quantitative Score for automated selection of optimal GAN hyper-parameters

We used T-PCAVR quantitative score to select our customized GAN with optimal hyper-parameters and enable the reproduction of results. Tab. 1 provides a comprehensive overview of the controlled parameters and their corresponding values, which were input into the T-PCAVR quantitative score. Subsequently, the T-PCAVR score determined the most optimal combination of these controlled parameter values for the GAN model. The selected GAN configuration includes 3 latent space variables randomly sampled from uniform distribution over half open interval [-1, +1). The Generator of GAN consists of 3 hidden layers with 200, 350 and 500 neurons respectively. The hidden layers of discriminator are exactly the same but in reverse order. Neurons of hidden layers are randomly initialized using Xavier Initialization. RMSProp algorithm is used as optimization function. Learning rate of 0.005 is used for both Generator and Discriminator. GAN is trained with the batch size of 64. To substantiate the efficacy of this scoring system in replicating comparable synthetic data in early human embryonic scRNA-seq dataset, as depicted in Fig. 6(right), supplementary experiments were conducted. The detailed outcomes of these experiments are concisely outlined in the sub-section 3.11. Additionally, the T-PCAVR score successfully applied to another limited dataset of mouse embryonic scRNA-seq data [34]. A summary of these results can be found in sub-section 3.12. However, T-PCAVR score have a limitation, detailed in sub-section 3.13

### 3.4 Evaluating GAN for Recovering High Order Relationship as Hidden Knowledge

We independently applied *ab-initio* knowledge discovery algorithm on real and synthetic datasets. The algorithm selected same combination of time-point and cluster numbers as reference in both datasets. The samples of E5 with 3 clusters is used to recover cell-lineage developmental story for both real and synthetic datasets.

**Table 2:**
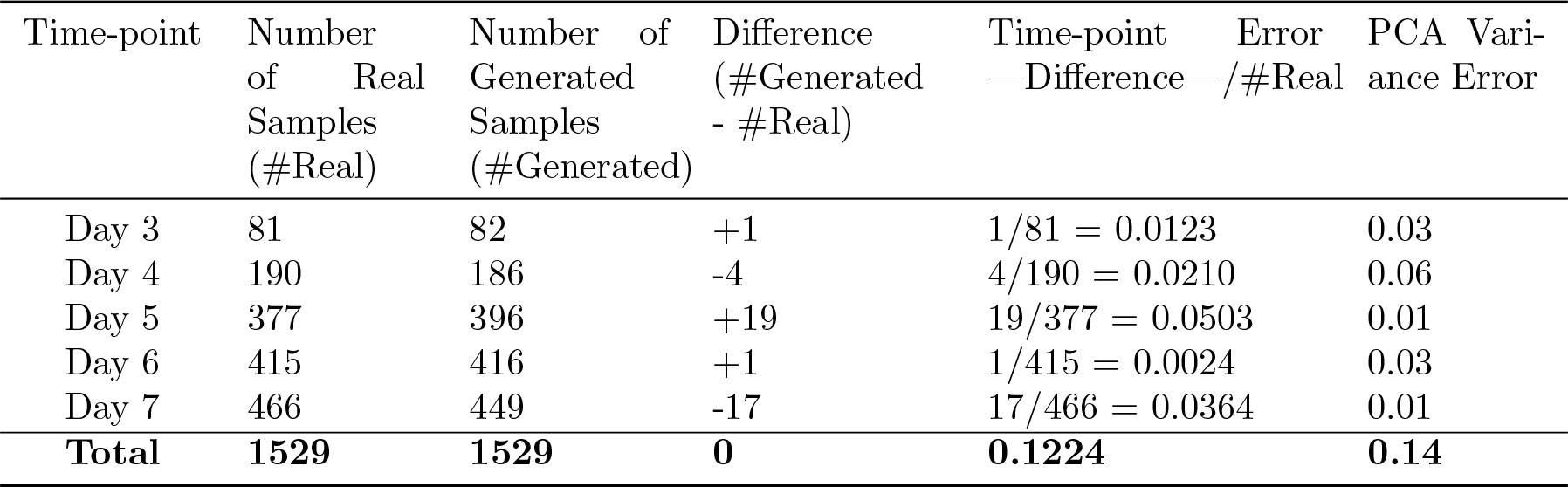
Quantitative summary of the samples in each time-point of real and generated datasets.

**Table 3:**
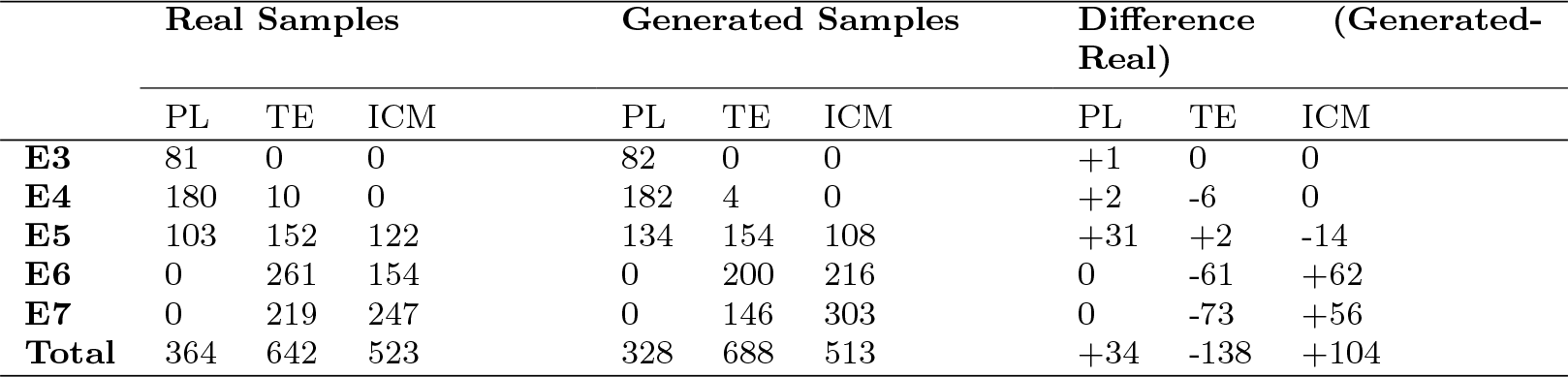
Distribution of real and generated samples for each time-point w.r.t lineages.

|  | Real Samples |  |  | Generated Samples |  |  | Difference (Generated-Real) |  |  |
| --- | --- | --- | --- | --- | --- | --- | --- | --- | --- |
|  | PL | TE | ICM | PL | TE | ICM | PL | TE | ICM |
| **E3** | 81 | 0 | 0 | 82 | 0 | 0 | +1 | 0 | 0 |
| **E4** | 180 | 10 | 0 | 182 | 4 | 0 | +2 | -6 | 0 |
| **E5** | 103 | 152 | 122 | 134 | 154 | 108 | +31 | +2 | -14 |
| **E6** | 0 | 261 | 154 | 0 | 200 | 216 | 0 | -61 | +62 |
| **E7** | 0 | 219 | 247 | 0 | 146 | 303 | 0 | -73 | +56 |
| **Total** | 364 | 642 | 523 | 328 | 688 | 513 | +34 | -138 | +104 |

*ab-initio* knowledge discovery algorithm provides us with two complete cell-lineage developmental stories from E3 to E7 for real and synthetic datasets. From the original work [33], there are three major cell types from E3 until E7. E3 and E4 samples consists of pre-lineage cells. E5 samples consists of pre-lineage, TE and inner cell mass (ICM) cells. Samples of E6 consists of TE and ICM cells while pre-lineage cells completely disappear. E7 samples also consist of TE and ICM cells only. Fig. 7 and Fig. 8 shows that cell-lineage developmental story of real and synthetic dataset are identical in terms of appearance and disappearance of cell-lineages at different time-points. These results are quantitatively summarized in Tab. 2. Fig. 9 and Tab. 3 provides a more comprehensive summary of the high order relationship of cell-lineage development in relation with each time-point. Each vertical stack in Fig. 9 is for one specific time-point. They are orderly arranged from left to right for E3 to E7 samples. The 3 different colors represent cell-lineages. Blue for pre-lineage (A), green for TE (C), and orange for ICM (B). Fig. 9 (right) is stack plot of generated samples. Fig. 9 (left) is stack plot of real samples. This Figure gives better comparative insight into the distribution of synthetic and real samples in different time-points with respect to cell-lineages.

**Fig. 7:**
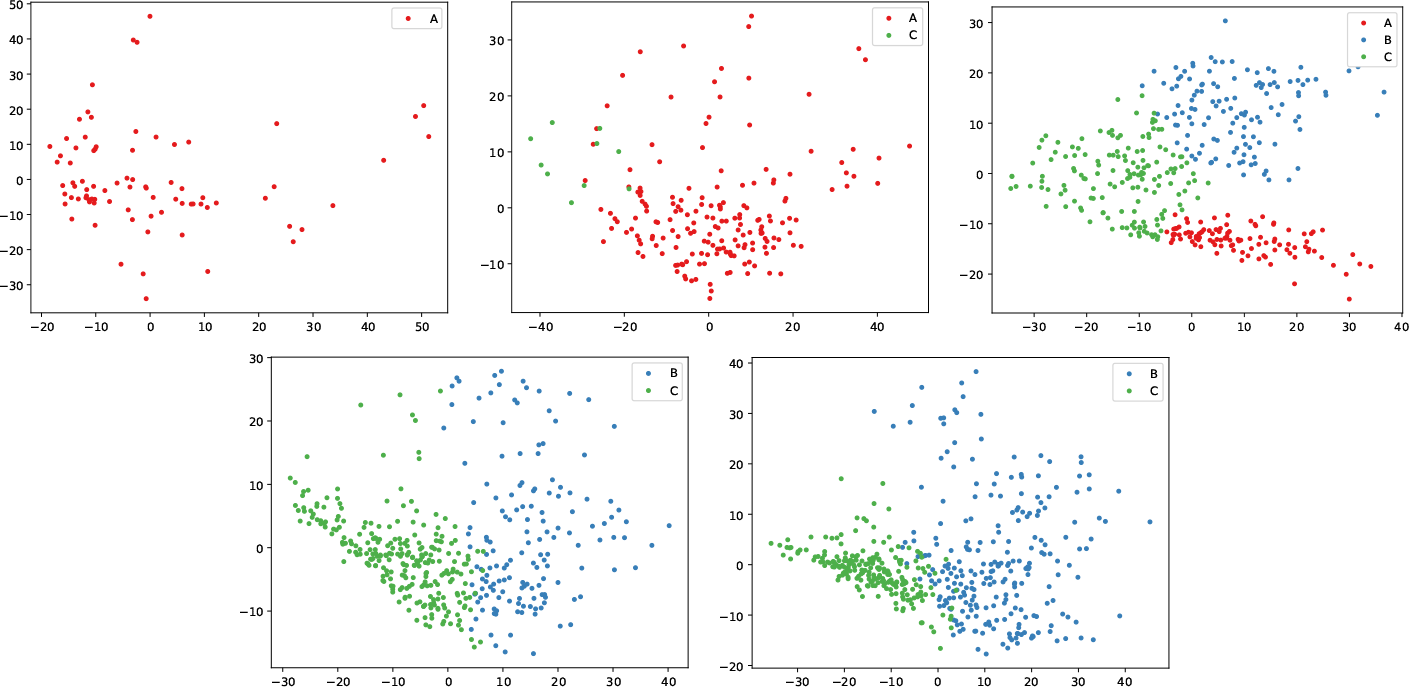
2D PC projection of 490 genes w.r.t cell-lineages in time-point E3(top left), E4 (top middle), E5 (top right), E6 (bottom left) and E7 (bottom right) for real 1529 samples

**Fig. 8:**
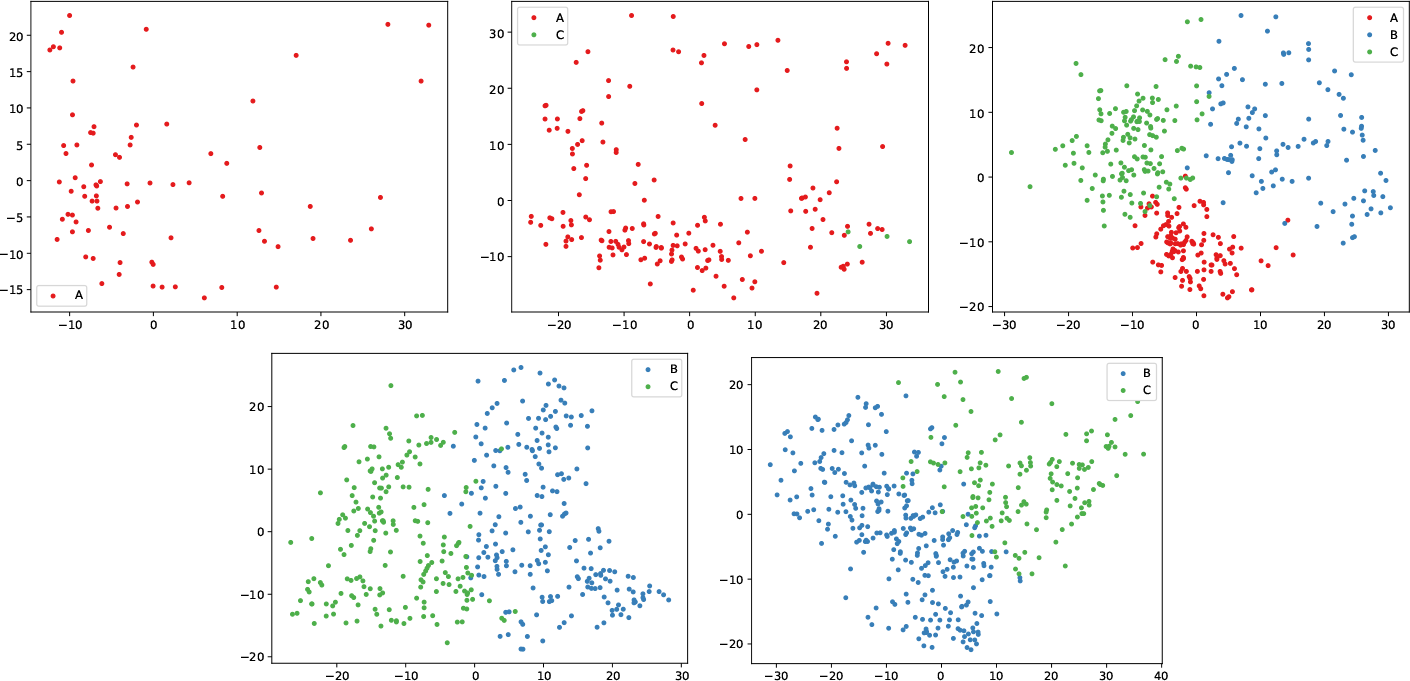
2D PC projection of 490 genes w.r.t cell-lineages in time-point E3(top left), E4 (top middle), E5 (top right), E6 (bottom left) and E7 (bottom right) for generated 1529 samples

**Fig. 9:**
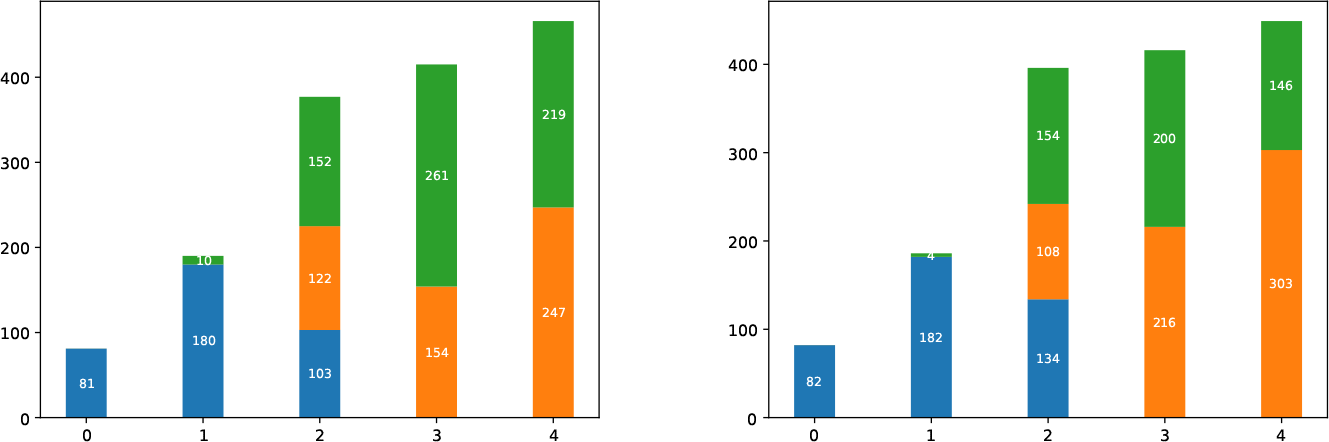
Stack plot of Real (left) vs Generated (Right) Cell developmental Story

### 3.5 Comparing our customized cwGAN with state-of-the-art cscGAN model on Small Dataset

To determine the efficacy of our optimally tuned customized cwGAN model, we conducted a comparative study with the state-of-the-art cscGAN model [18]. Both models used time-points as the conditional input variable. The cscGAN model was subjected to 10,000 training epochs, beyond which the model failed to exhibit any further improvement. Additionally, we repeated the experiment of training cscGAN model for six individual runs and reported the top two results in this section.

As explained earlier, we did not rely on conditional input to decide the time-point label for the samples generated with cwGAN model. Rather, we trained an svm classifier on samples of real human embryonic scRNA-seq dataset and then predicted the samples of generated dataset for robust downstream analysis. In the same way, we identified the time-point labels for the cscGAN generated datasets. However, the results were unsatisfactory as shown in Fig. 10 (column 2). In this regard, we relaxed the condition and kept the conditional input as time-point label as shown in Fig. 10 (column 1). Next, we applied the exhaustive search algorithm on the cscGAN generated datasets. The algorithm identifed E7 with six clusters and E7 with seven clusters for the two datasets. A comparative analysis of the results obtained from the exhaustive search algorithm on the real human embryonic dataset with cscGAN generated datasets exposed inconsistencies. These inconsistencies from cscGAN generated datasets. To address this, we relaxed the condition for recovering the cell lineage developmental story in an ab-initio manner for cscGAN generated datasets. Instead, we manually select the sample of E5 with three clusters as reference. This reference choice, identified by exhaustive search algorithm, had previously yielded anticipated results in both the real and cwGAN generated dataset. Despite this additional relaxation, the cscGAN generated datasets failed to preserve the hidden high-order relationships within the cell lineage developmental story. The representation of the cell lineage developmental story for the two cscGAN-generated datasets is presented in Fig. 11, using a stackplot. From prior biological knowledge, pre-lineage cell types can not exist in E6 and E7. Conversely, inner cell mass (ICM) cannot exist in E4, and both ICM and trophectoderm (TE) cannot exist in E3. In both cscGAN-generated datasets, should one interpret the cell type (depicted in blue in Fig. 11) as pre-lineage, the dominant presence of pre-lineage in E6 and E7 contradicts established biological knowledge, indicating a failure of the cscGAN model to preserve hidden knowledge. Similarly, interpreting the dominant cell type as TE or ICM leads to a cell developmental story that lacks logical coherence from a biological perspective.

**Fig. 10:**
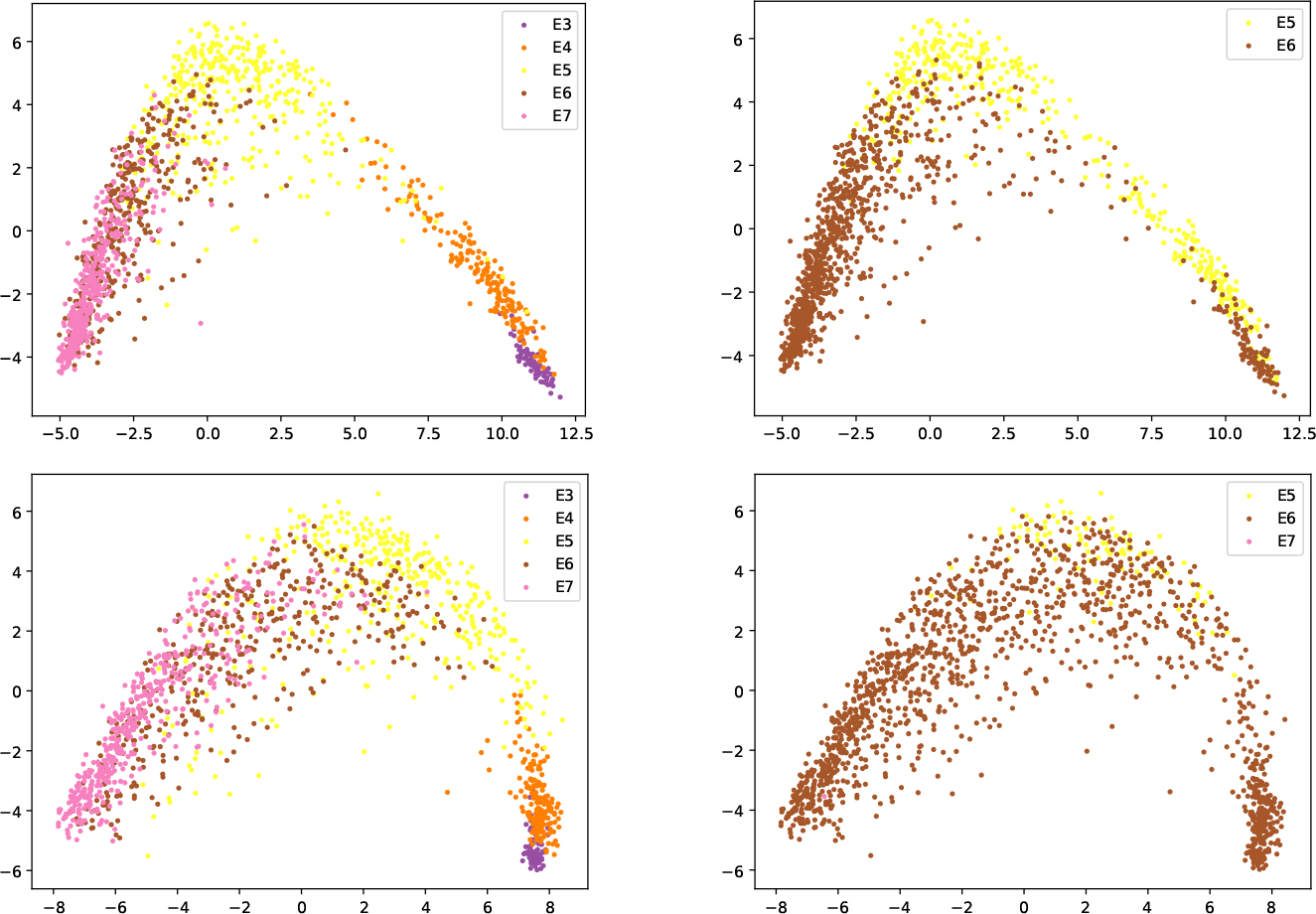
Top two 1529 samples datasets (top and bottom) generated by cscGAN trained with conditional label as cell sequencing time-points for Early Human Embryonic scRNA-seq developmental dataset. The 2D PC plot is labelled with conditional labels (left) and svm identified labels (right)

**Fig. 11:**
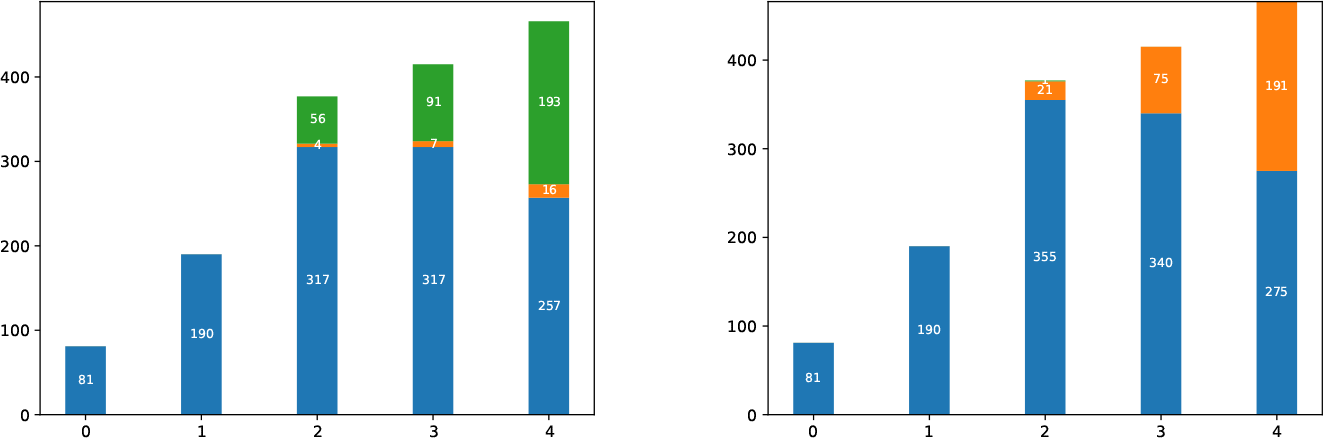
Cell lineage developmental story w.r.t embryonic time-points for the top two generated datasets (left and right) by cscGAN

It is noteworthy that we trained the cscGAN model on two versions of the real human embryonic datasets. One comprising the complete set of real human embryonic genes (26178 genes) and the other consisting of only 490 highly variable genes set.

It is crucial to emphasize that our comprehensive analysis and subsequent reporting exclusively consider the optimal results derived from training with the complete real human embryonic dataset encompassing all genes.

It is noteworthy, that we attempted to train cscGAN not only on the real human embryonics with all genes (26178) but also on 490 highly variable genes as well. Nevertheless, we only included the best results achieved by training cscGAN with all genes.

### 3.6 Timepont LSI Scheme Shows the Preservation of Known Information

Fig. 12 shows the impact of small alterations in GAN’s latent space variable onto the time-point semantic information of the dataset. This result verifies that GAN captured time-point as known information in its latent space. The time-points of synthetic samples are represented with colors of red (E3), teal (E4), light green (E5), dark blue (E6) and black (E7).

**Fig. 12:**
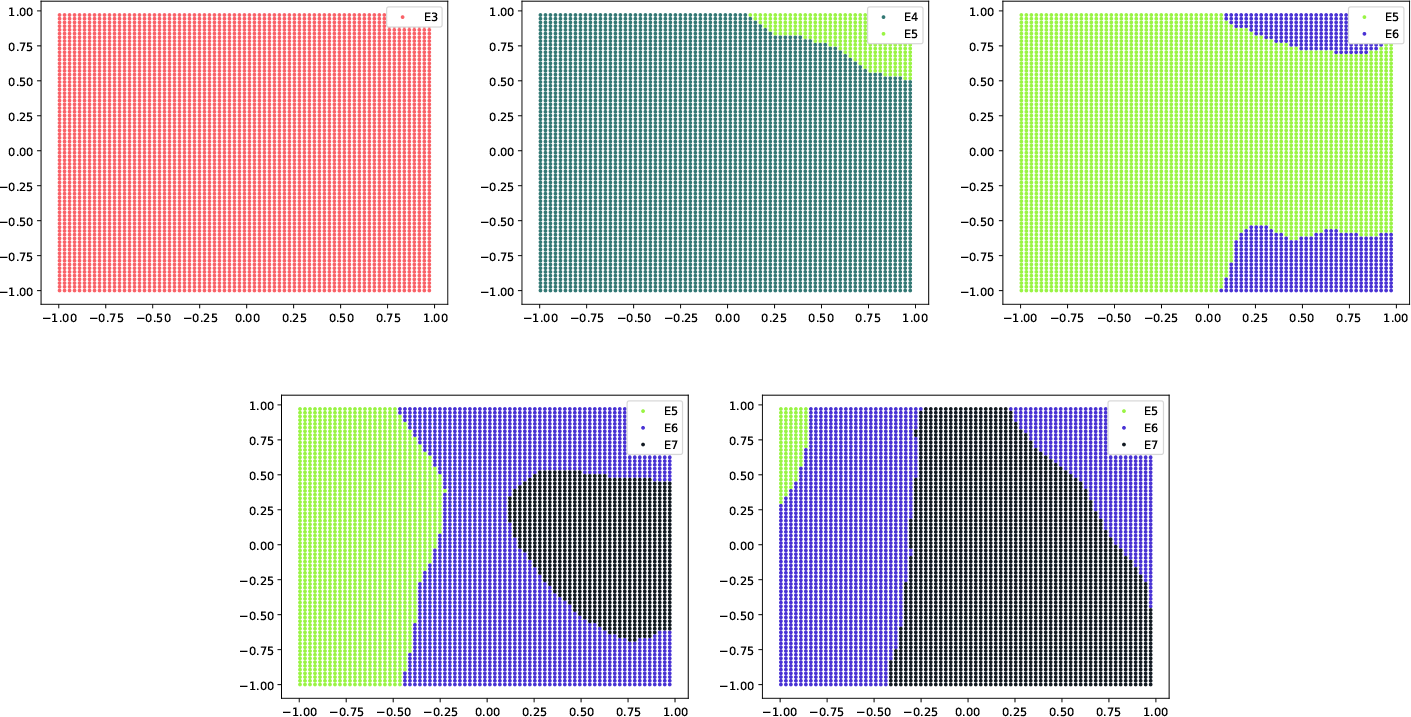
Latent space interpretation for generated samples in terms of time-points

### 3.7 Efficacy of Identified Time-points Compared to Conditional Input

As mentioned, we used conditional input as regularization step for the better training of GAN. Conventionally conditional input alone decides the label of generated sample [18, 35]. However, in Fig. 12, we observe that time-point as conditional input does not necessarily correspond to the identified time-point for the same synthetic sample. This aspect has been briefly discussed in the prior research [43]. To show the more accurate time-point among the two, we map the first 2 PC components of generated synthetic samples from Fig. 12 onto the 2D PC plot of real 1529 dataset in Fig. 6(left). Time-points of real dataset is used as reference. After mapping, we evaluate if time-points as conditional input have better overlap with the defined reference compared to identified time-points. It is evident from Fig. 13 that identified time-points are logically mapped better than conditional inputs.

**Fig. 13:**
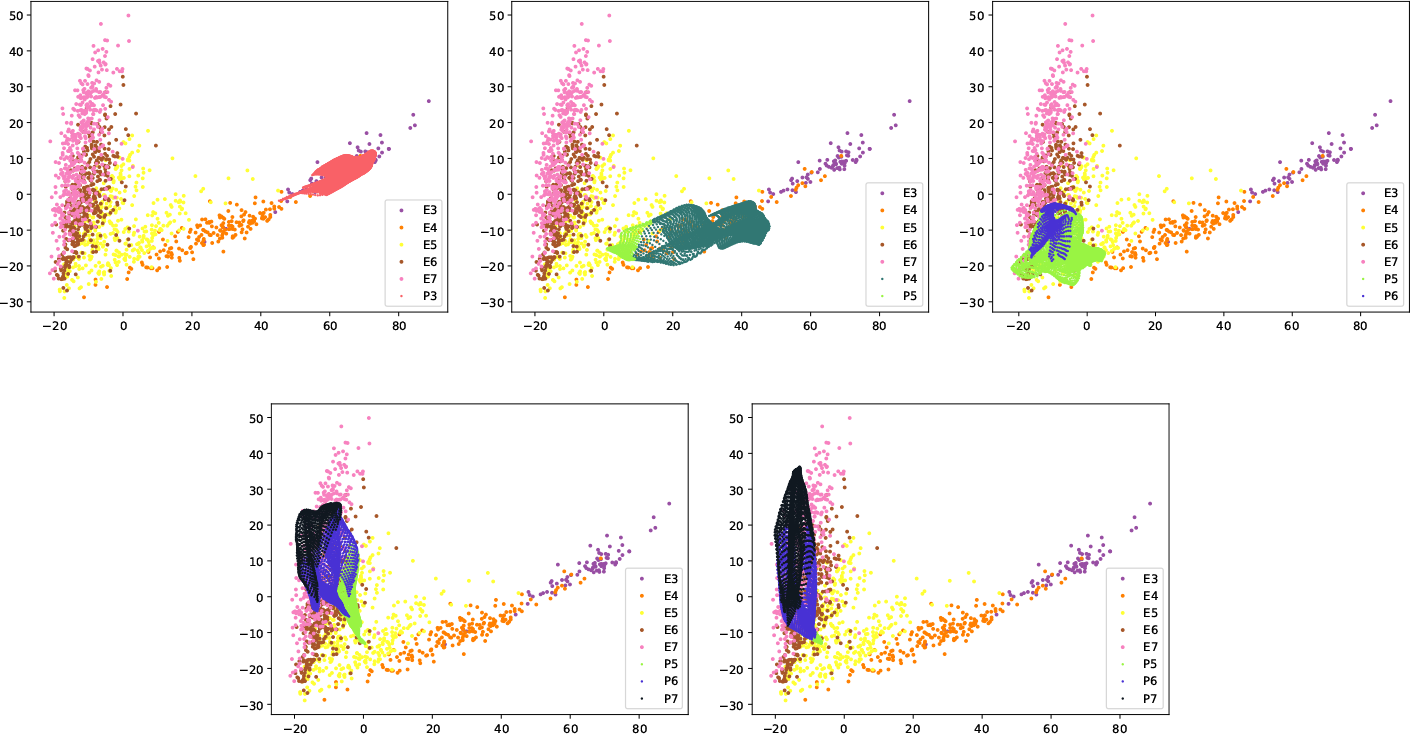
Mapping generated samples onto real 1529 samples labeled for time-points

### 3.8 Lineage LSI Scheme Shows the Preservation of Hidden Knowledge

Fig. 14 demonstrates the nuanced relationship between the latent space variables of GAN and cell-lineage developmental information. This result shows that GAN successfully captured hidden knowedge as high-order relation of cell-lineage by encoding cell-lineage as unknown semantic in the latent space.

**Fig. 14:**
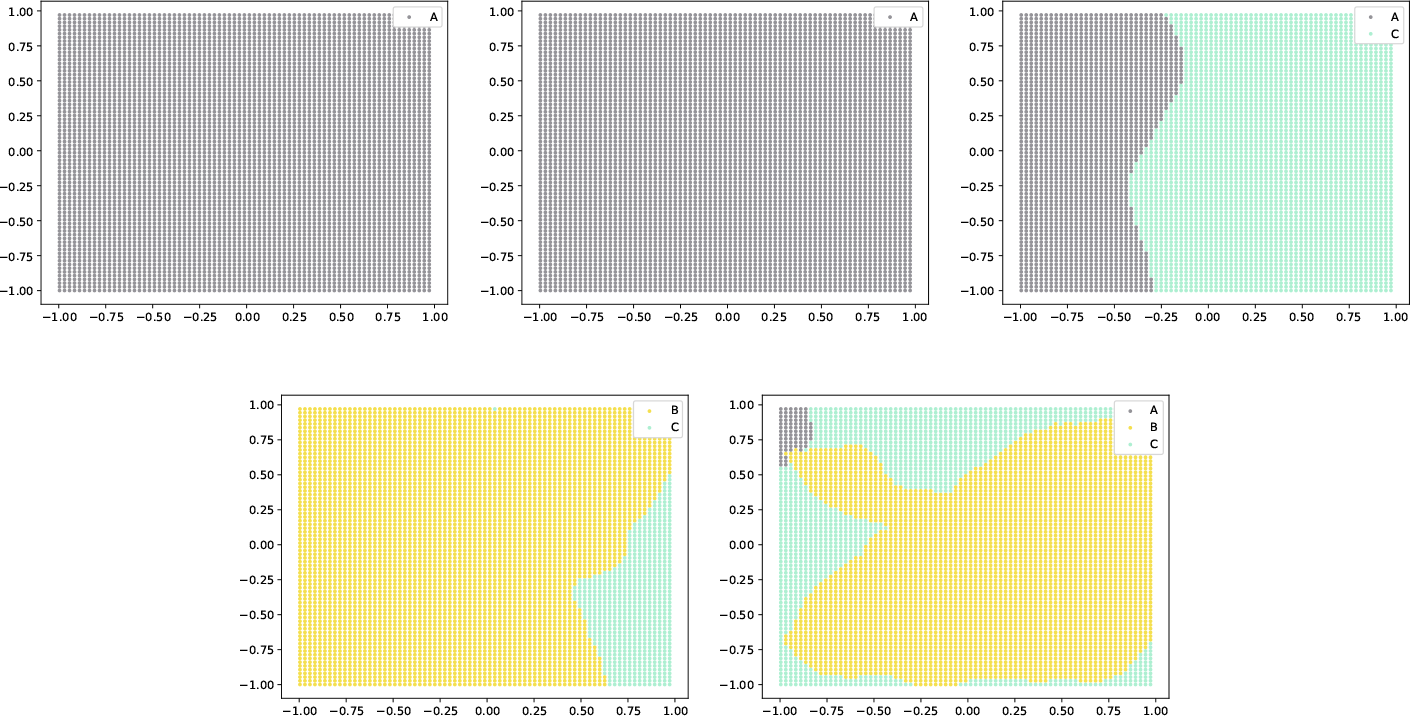
Latent space interpretation for generated samples in terms of cell-lineages

### 3.9 Efficacy of Identified Lineages Compared to Conditional Input

In Fig. 15, we map the first 2 PC components of generated synthetic samples from Fig. 14 onto the real 1529 dataset PC plot in Fig. 6(left). Cell-lineages of real dataset is used as reference. We observe significant overlap of corresponding cell-lineages between synthetic samples and real 1529 dataset.

**Fig. 15:**
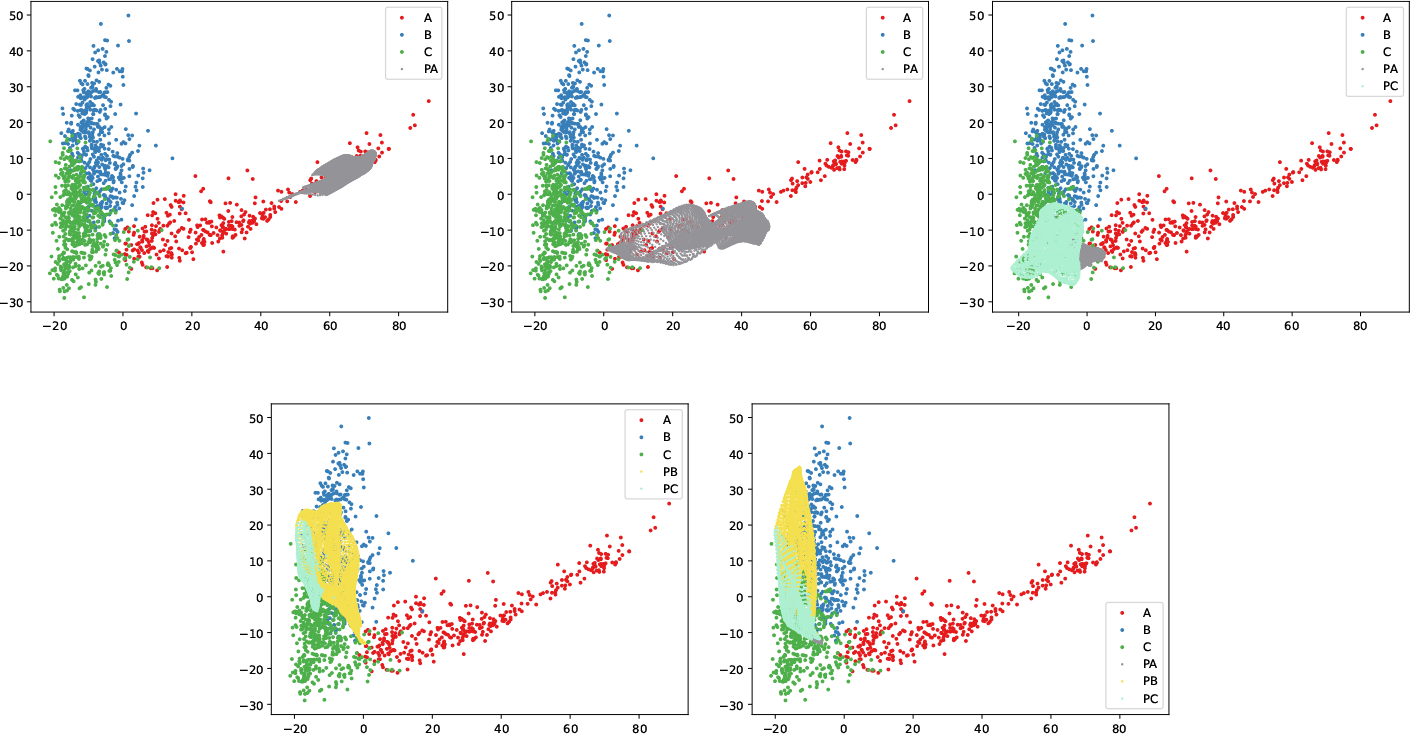
Mapping generated samples onto real 1529 samples labeled for cell-lineages

### 3.10 Comprehensive Database to Sample Precise Time-Point and Cell-Lineage Cells and Identifying Anomalies

For deeper insights, we study the relationship of latent space variables with time-points and cell-lineages combined. In the combined scheme of experiment, cell-lineages are represented with filled small sized circle as A (pre-lineage), filled medium sized triangle as B (ICM) and filled large size star as C (TE). Whereas, five time-points are represented with red (E3), teal (E4), light green (E5), blue (E6) and black (E7) colors.

Using this scheme, we created a database of approximately two million unique input combinations. Each combination consists of three latent space variables and one conditional variable. All the unique combinations for three latent space variables are acquired from sets of Eq. 1, Eq. 2 and Eq. 3. With the help of this database, we can identify the exact number of unique input combinations to get a sample of specific time-point and cell-lineage. For example, we exactly know the 1,99,665 unique input combinations that provide us with pre-lineage samples of E5 or 2,38,355 unique input combinations that gives us TE samples of E6 or 2,77,102 unique input combinations that results in ICM samples of E7, etc.

Moreover, using cell-lineage development of real 1529 dataset in Fig. 7 as reference story, we can identify all unique input combination that results in conflicting time-point and cell-lineage. We searched the entire database of approximately two million instances and found some unique input combinations that resulted in pre-lineage samples of E6 and E7. This is depicted in Fig. 16b. where red cross represents contradicting time-point and cell-lineage and green filled circle represents those inline with reference story. 2D Interpretation plot in Fig. 16a shows some samples of E6 that belong to prelineage. These samples are represented with small blue filled circles. Since the samples contradict with our reference story therefore the input combinations that generated such synthetic samples are marked as outliers.

**Fig. 16:**
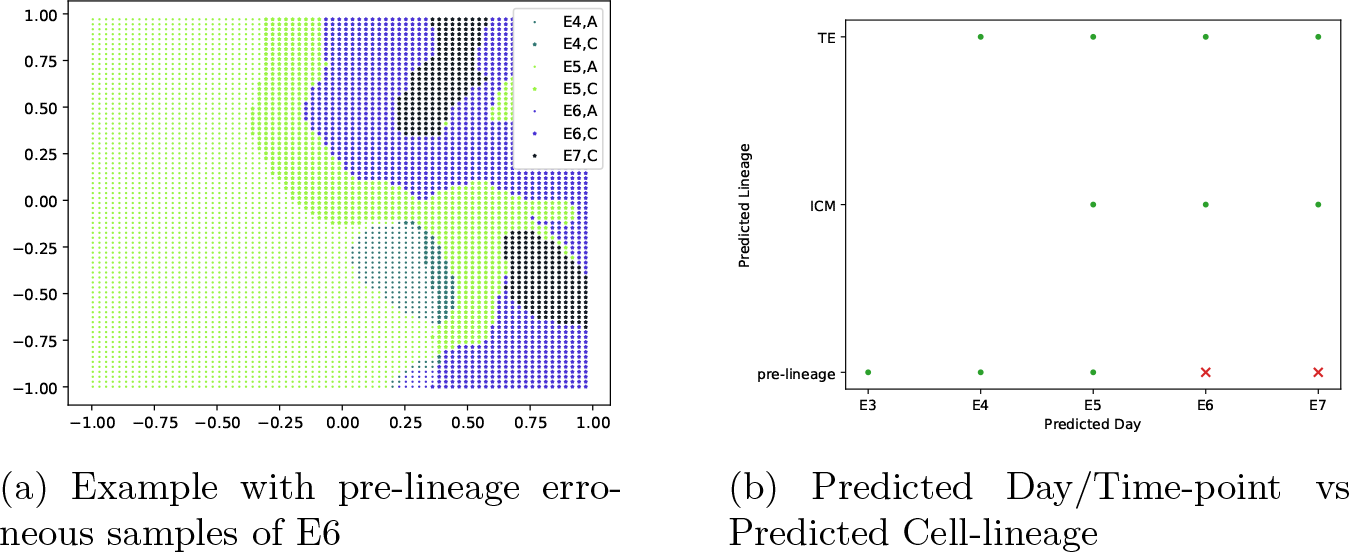
Combined interpretation scheme w.r.t time-points and cell-lineages identify anomalies

### 3.11 Reproducing GAN-generated Data for Early Human Embryonic scRNA-seq Data with T-PCAVR Score

Training GANs is widely acknowledged as a formidable challenge within the research community. This challenge becomes significantly more daunting when attempting to train a GAN using a dataset of very limited size. Moreover, replicating the results of GANs trained on such small datasets poses an even greater challenge. Existing literature has shed light on the limitations of current quantitative scores, such as Inception Score, particularly when applied to limited datasets with complex properties. In the context of our research, the dataset employed is exceptionally small, comprising only 1529 samples. The dataset is characterized by complex biological phenomenon of early human embryonic cell-lineage differentiation over a period of 5 days (day3 to day 7). Quantitative values like loss values of Generator and Discriminator and Inception Score failed to reproduce GAN results. Therefore, we formulated a T-PCAVR Quantitative score in order to reproduce GAN results. An example of the reproduced results compare to originally generated synthetic dataset is shown in Fig. 17 and Fig. 18

**Fig. 17:**
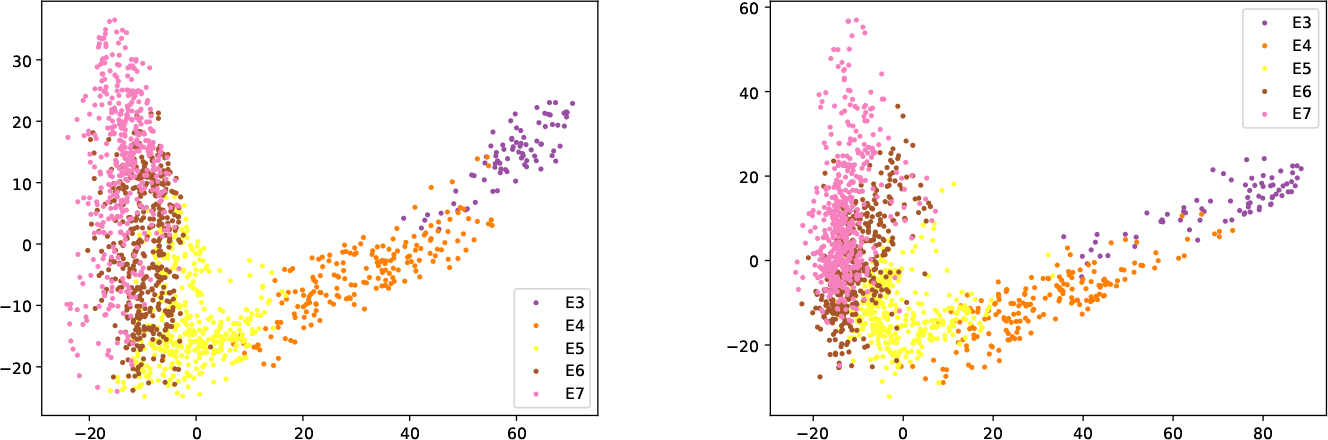
2d PC projection of 490 genes with 1529 samples for Generated (left) and Reproduced Generated (right) datasets

**Fig. 18:**
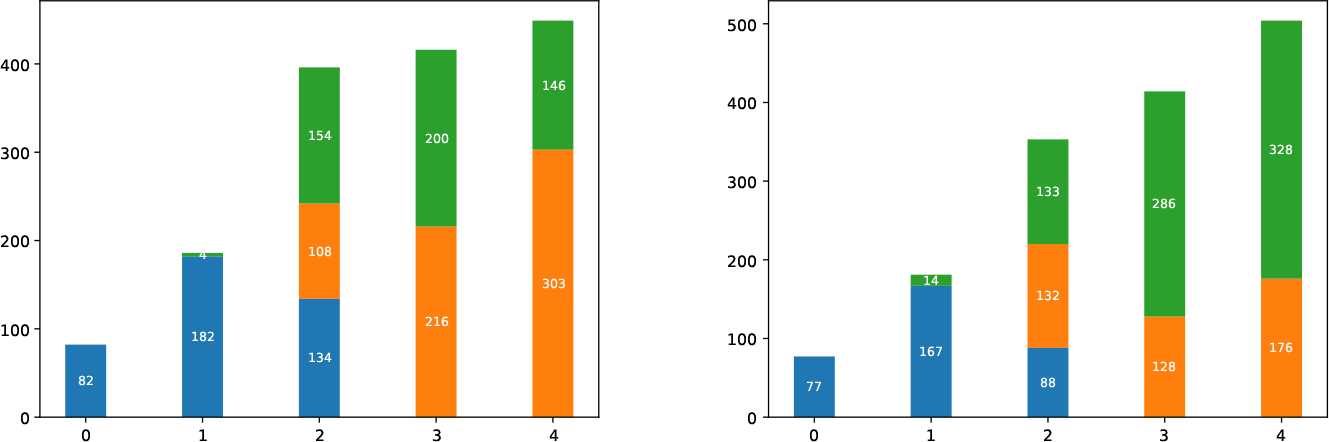
Stack plot of Generated Cell developmental Story (left) vs Generated Cell developmental Story reproduced with T-PCAVR Score (right)

### 3.12 GAN-generated Data for Early Mouse Embryonic Data with T-PCAVR Score

To assess the effectiveness of our experimental setup, where we recover hidden knowledge by selecting an optimally trained GAN using the T-PCAVR quantitative score, we endeavored to generate synthetic data for an early mouse embryonic scRNA-ses dataset [34]. The early mouse embryonic scRNA-seq dataset consists of 1724 single cells sequenced in embryonic timepoint E5.25, E5.5, E6.25 and E6.5. There are 500 highly differenciated genes for each cell. Notably, the SVM with RBF kernel failed to properly learn the time-points from the real dataset. When SVM was trained on the sample’s time-points, it inaccurately predicted all generated samples to be at time-point E6.5 of mouse embryonic development. Consequently, we substituted SVM with a Random Decision Forest [40] to learn the time-points from the real data, which resulted in accurate predictions for the time-points of the generated dataset. Fig. 19 shows the real (left) and synthetic (right) mouse embryonic scRNA-seq datasets labelled for time-points. Fig. 20 shows the cell-lineage developmental story recovered by *ab-initio* method on real early mouse embryonic scRNA-seq dataset. Fig. 21 shows the cell-lineage developmental story recovered by *ab-initio* method on synthetic early mouse embryonic scRNA-seq dataset. Fig. 22 illustrates the cell-lineage types (stacks within each vertical bar) corresponding to each time-point (vertical bars). Notably, the real dataset, when labeled with time-points, lacks distinct segregation as shown in Fig. 19(left). Consequently, the comparison between the real and synthetic datasets is evident in the context of cell-lineage types, as demonstrated in Fig. 20, Fig. 21 and Fig. 22.

**Fig. 19:**
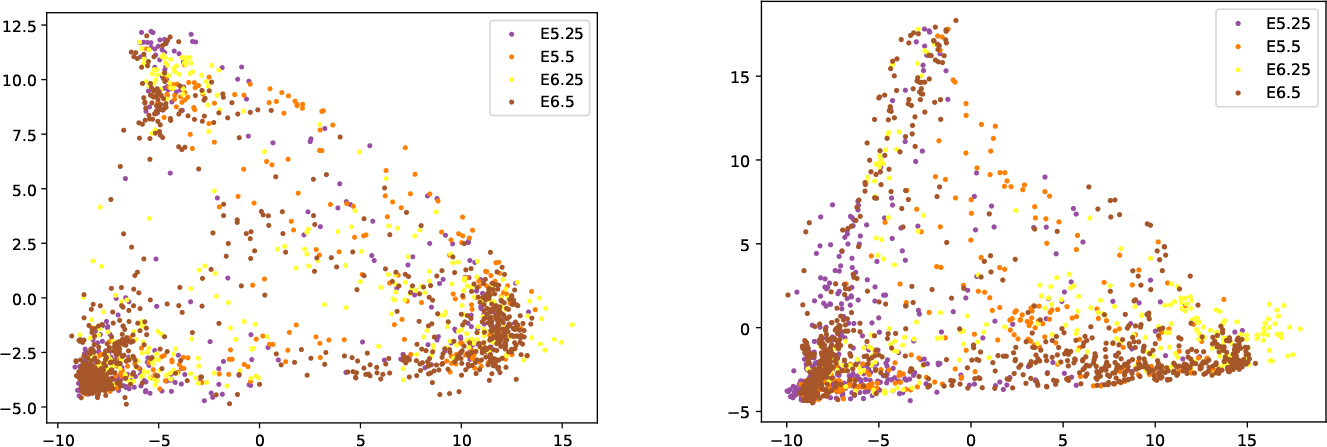
2d PC projection of 500 genes with 1724 samples for real (left) and generated (right) early mouse embryonic datasets

**Fig. 20:**
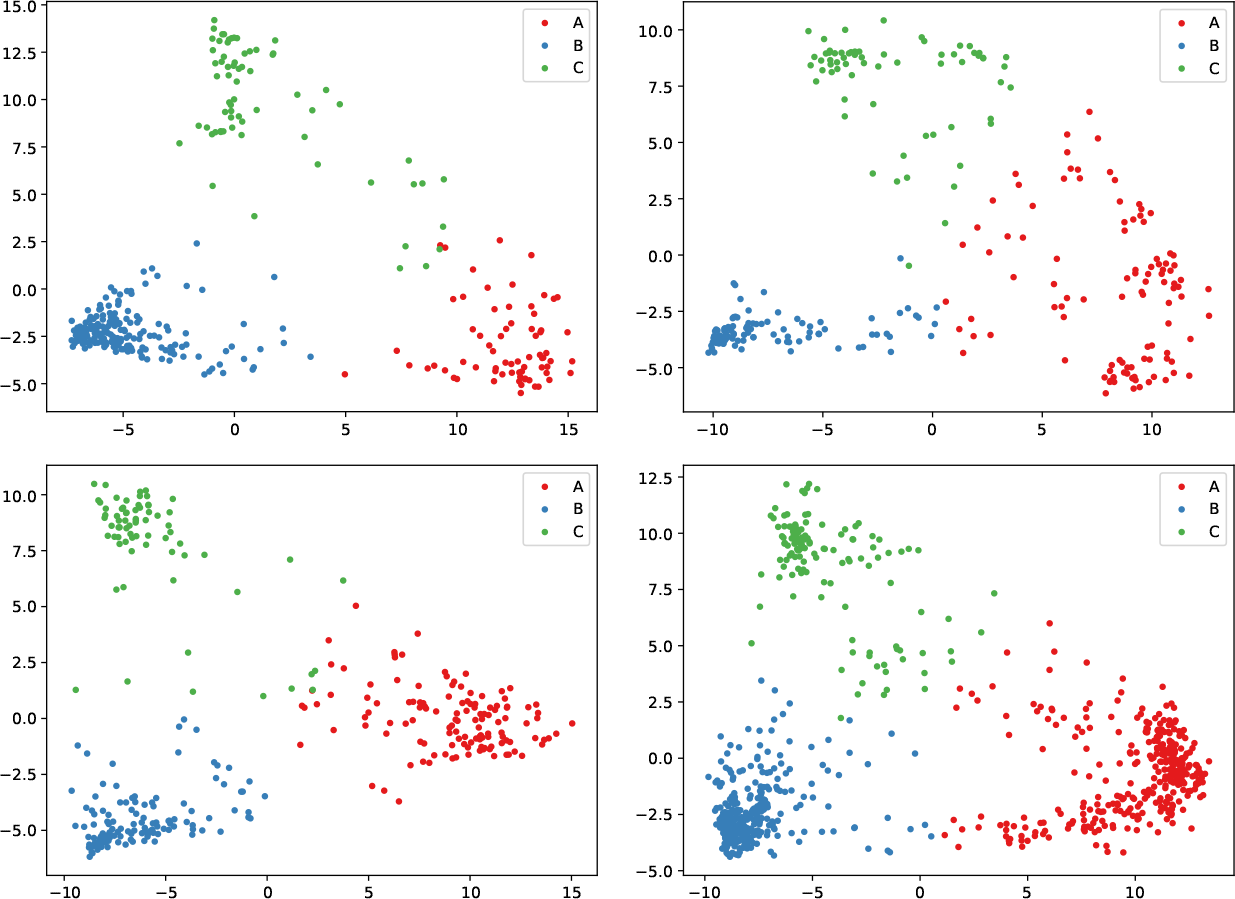
2D PC projection of 500 genes w.r.t cell-lineages in time-point E5.25(top left), E5.5(top right), E6.25(bottom left) and E6.5(bottom right) for real 1724 samples

**Fig. 21:**
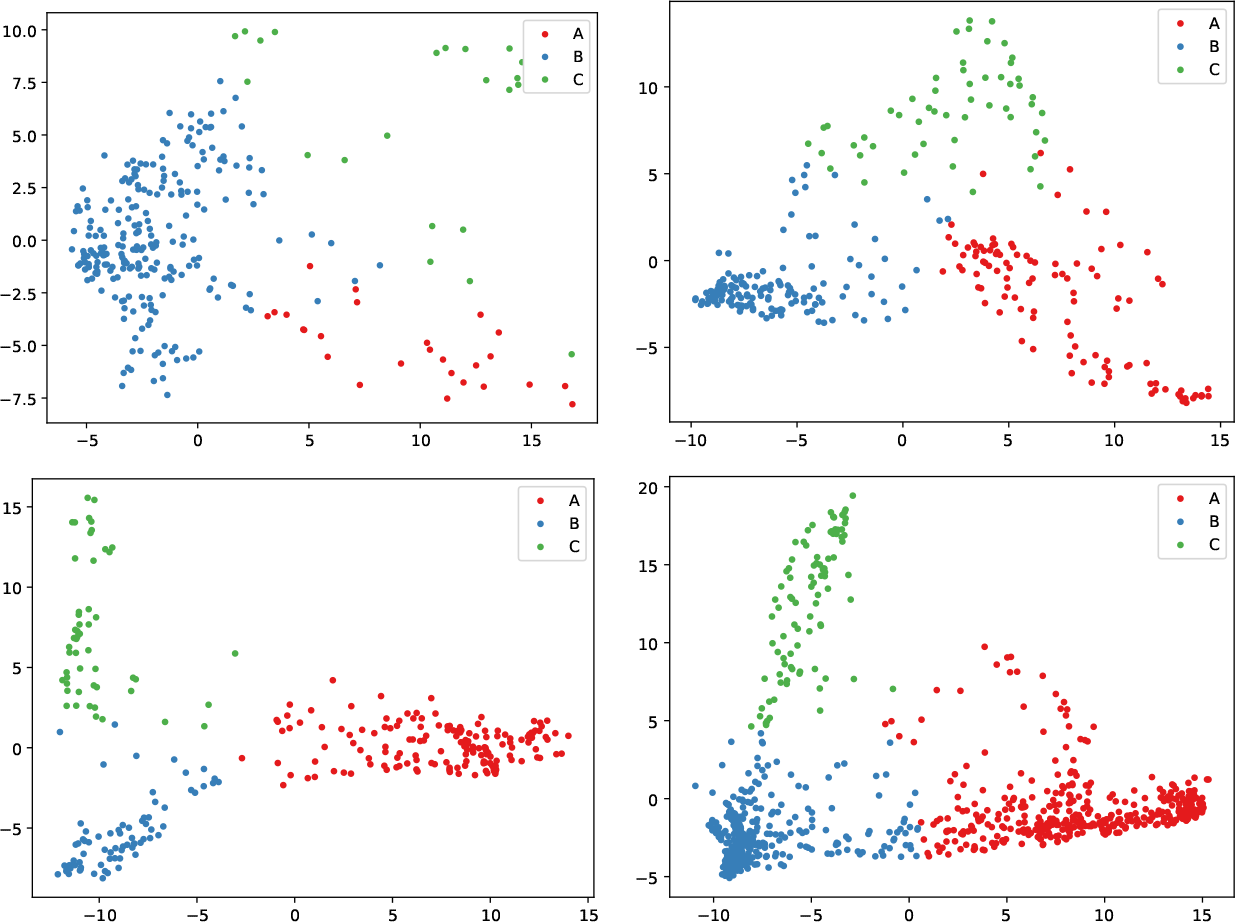
2D PC projection of 500 genes w.r.t cell-lineages in time-point E5.25(top left), E5.5(top right), E6.25(bottom left) and E6.5(bottom right) for generated 1724 samples

**Fig. 22:**
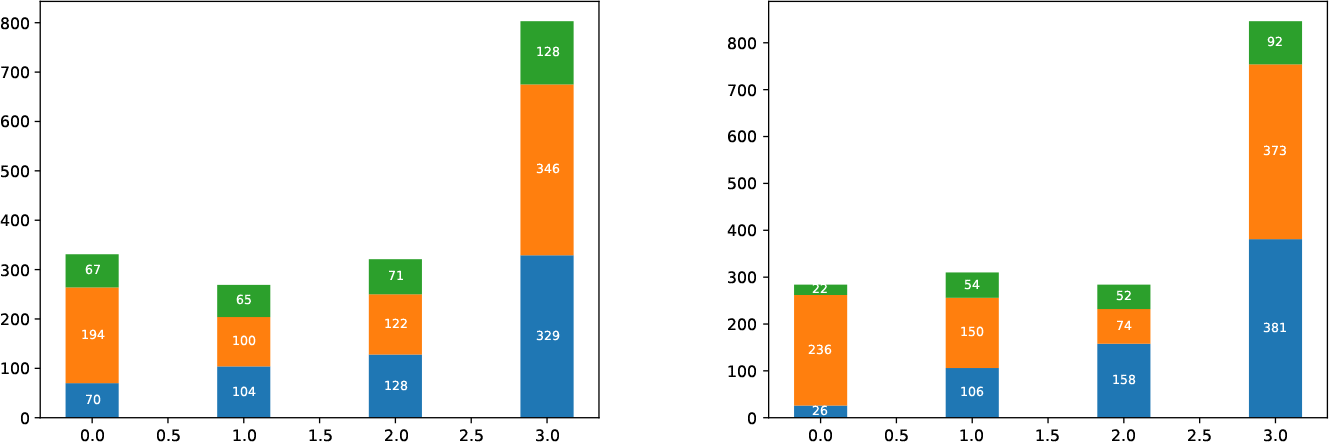
Stack plot of Real (left) and Synthetic (right) Cell developmental Story

### 3.13 Limitation of T-PCAVR Score

One of the major drawbacks of the T-PCAVR Score is its inability to deter the selection of suboptimal hyper-parameters when the GAN is susceptible to overfitting. We conducted further experiments into one of our suboptimally trained GAN model suffering from overfitting. We found that the overfitting was primarily manifested due to the cell-lineage types within the data. An instance where the T-PCAVR Score incorrectly identified, configuration of GAN model suffering from overfitting caused by cell-lineage types, is illustrated in Fig. 23 and Fig. 24. Fig. 24 provides a detailed illustration demonstrating that the number of samples at each time-point in the synthetic dataset is relatively consistent with that of the real dataset. This phenomenon can be observed by comparing the height of the bars in the two sub-plots of Fig. 24. However, when viewed from the perspective of cell-lineage types, it is evident that there is a marked over-representation of the cell-lineage type represented in green (TE) in Fig. 24. This over-representation drastically alters the lineage developmental story when compared to prior knowledge.

**Fig. 23:**
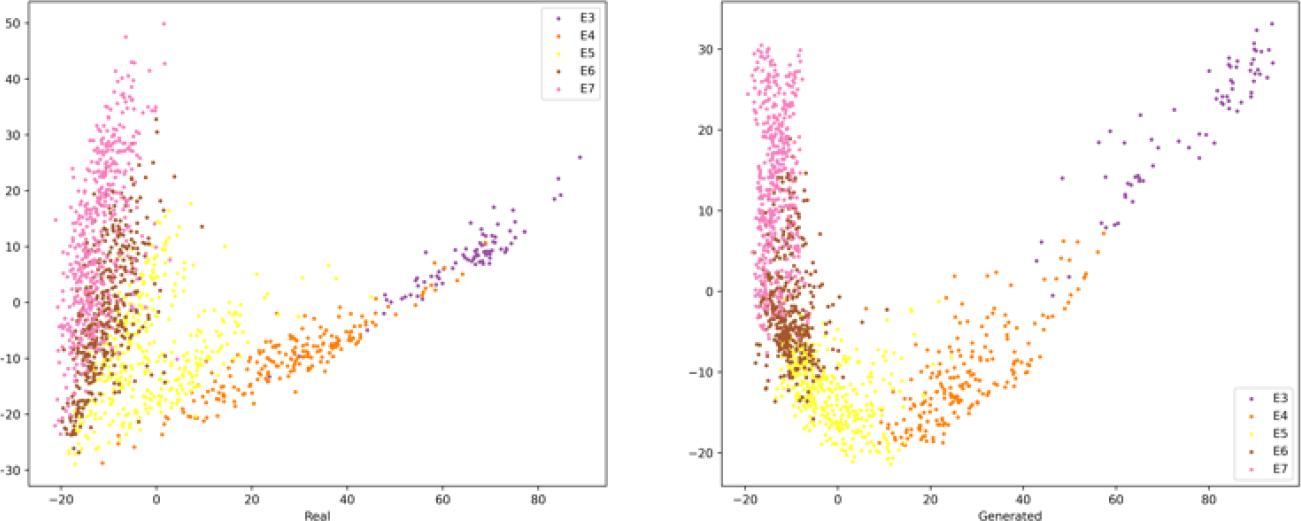
An example of Optimal GAN model identified by my T-PCAVR score. Left subplot is 2D PC plot of real dataset and Right subplot is 2D PC plot of synthetic dataset

**Fig. 24:**
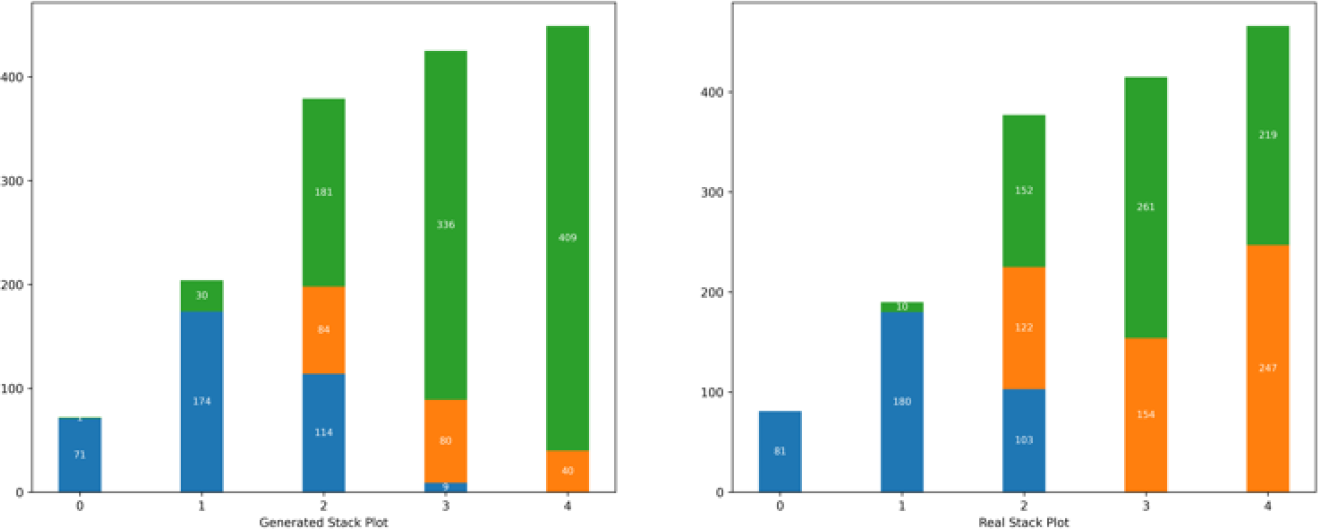
Stackplot to highlight the overfitting due to cell-lineage types

Given that the primary goal of employing the *ab-initio* knowledge discovery method is to evaluate the GAN’s ability to capture and preserve the lineage developmental process as hidden knowledge, therefore utilizing lineages to enhance the T-PCAVR Score is not a feasible solution.

## 4 Discussion and Conclusion

This study recovered hidden knowledge from real and synthetic GAN-generated samples in *ab-initio* manner. Furthermore, the research work evaluated the quality of the custom GAN-generated samples and conducted a comparative analysis with state-of-the-art cscGAN model. The cell-lineage developmental story was used as hidden knowledge, to be recovered in *ab-initio* manner, in order to evaluate the performance of GAN model. A quantitative error score was formulated to reproduce optimally tuned GAN model. Additionally, LSI schemes were designed to investigate how GAN-generated data captured high-order relations from real data. The LSI schemes were used to understand the encoding of known (time-points) and unknown (cell-lineages) semantics in latent space of GAN. The combined LSI scheme of cell-lineage and time-point was further used to identify anomalies and sample desired cell of specific time-point and cell-lineage.

Experimental results show that the quality of synthetic samples was more superior than cscGAN model on a limited dataset with complex properties. T-PCAVR quantitative error score successfully reproduced quality synthetic data for human and mouse embryonic datasets with optimally tuned GAN configurations. *ab-initio* knowledge discovery method recovered hidden knowledge of cell-lineage developmental story from GAN-generated synthetic dataset. Thus providing a more objective evaluation about the performance of our custom GAN model. The hidden knowledge recovered from GAN-generated synthetic dataset was identical to the hidden knowledge recovered from real dataset. Additional results demonstrate that LSI schemes provide a visual representation about how GAN preserved the hidden knowledge by successfully capturing lineage developmental story as unknown semantic in the latent space. Time-point Cell-lineage LSI scheme identified anomalies after verifying with prior knowledge and enabled the sampling of cells with specific time-point and cell-lineage.

In future, we intend to further explore latent space interpretation to interpolate data between two time-points or more generally when the sampling of real data is sparse. Furthermore, our future research agenda includes the presentation of real-world case studies, intended to inspire scientists across diverse disciplines to explore the research domain of ‘Knowledge Discovery from Synthetic Data’. Additionally, we plan to explore disentanglement of latent space variables coupled with large scale generated dataset for a more detailed timeline of lineage differentiation.

The limitation of this study is primarily the inability of T-PCAVR quantitative score to ignore the overfitting due to cell-lineages, explained in sub-section 3.13. Second limitation of this work is the assumption for *ab-initio* knowledge discovery method that samples of at least one time-point have representations for all cell-lineages.

In summary, discovery learning, using ML methods to extract scientific knowledge from real observations, is a promising research avenue. Notably, DGMs like GANs have shown promise in bridging the gap between real and synthetic samples. While many studies emphasize the utility of synthetic data for diverse applications, our research addresses less-explored yet crucial aspects in GAN-related studies. This research work underscores the vital need to assess DGM models, especially GANs, on limited, complex datasets and compare them with state-of-the-art model. Moreover, the inadequacy of current quantitative metrics for reproducing optimal GAN models, particularly on limited datasets, poses a significant challenge. Addressing this issue requires the development of data-specific quantitative metrics. Additionally, the lack of interpretability between the latent space of GANs and essential data semantics is a pressing concern. To address this gap, intuitive visual frameworks are necessary, establishing clear connections between the latent space of GANs and the important semantics with the dataset. Our study presents a systematic effort for answering some of the basic compelling questions using biological dataset. We believe our research can inspire scholars across disciplines to comprehend the complexities of synthesizing limited datasets with DGM models, tackle the challenges in reproducing optimal DGM models, adopt objective evaluations for data-specific hidden properties, and gain insights into how DGM models encode crucial data semantics.

## 5 List of Abbrevations

scRNA-seq: Single Cell RNA Sequencing
ML: Machine Learning
GAN: Generative Adversarial Networks
DGM: Deep Generative Models
SVM: Support Vector Machines
RBF: Radial Basis Function
RDF: Random Decision Forests
cscGAN: Conditional Single Cell Generative Adversarial Networks
cwGAN: Conditional Wasserstein Generative Adversarial Networks with Gradient Penalty using Label Smoothing
T-PCAVR: Time-point Principal Component Analysis Variance Ratio
LSI: Latent Space Interpretation
TE: Trophectoderm
PL: Pre Lineage
ICM: Inner Cell Mass
RMSProp: Root Mean Square Propagation
ARS: Adjusted Reliability Score
OVR: One vs Rest
PCA: Principal Component Analysis
E3: Embryonic Day 3
E4: Embryonic Day 4
E5: Embryonic Day 5
E6: Embryonic Day 6
E7: Embryonic Day 7
E5.25: Embryonic Day 5.25
E5.5: Embryonic Day 5.5
E6.25: Embryonic Day 6.25
E6.5: Embryonic Day 6.5

## Declarations

### Ethics approval and consent to participate

No ethics approval was required for the study.

### Availability of data and materials

The original scRNA-seq data of early human embryonic cells and corresponding ERCC spike-in reference data can be found at E-MTAB-3929

The original scRNA-seq data of early mouse embryonic cells and corresponding ERCC spike-in reference data can be found at (GEO: GSE109071).

The pseudocode for the T-PCAVR quantitative score, mathematical formulations and schematic structure of LSI scheme along with the detailed architecture of the customized cwGAN and its corresponding hyper-parameters, are comprehensively outlined in the Methods and Results section.

The generated synthetic datasets for both the early human embryonic scRNA-seq dataset and the early mouse embryonic scRNA-seq dataset are made available alongside this manuscript.

### Consent for publication

Not Applicable

### Competing interests

The author declares no competing interests.

### Funding

This work was support in part by the NSFC grant 62250005 and the Chinese Scholarship Council (CSC). The authors extend special gratitude to CSC for their invaluable support, without which this research would not have been possible.

### Authors’ contributions

## Notes

### Competing Interest Statement

The authors have declared no competing interest.

